# An intrinsically disordered transcription activation domain alters the DNA binding affinity and specificity of NFκB p50/RelA

**DOI:** 10.1101/2022.04.11.487922

**Authors:** Hannah E.R. Baughman, Dominic Narang, Wei Chen, Amalia C. Villagrán Suárez, Joan Lee, Maxwell Bachochin, Tristan R. Gunther, Peter G. Wolynes, Elizabeth A. Komives

## Abstract

Many transcription factors contain intrinsically disordered transcription activation domains (TADs), which mediate interactions with co-activators to activate transcription. Historically, DNA-binding domains and TADs have been considered as modular units, but recent studies have shown that TADs can influence DNA binding. We biophysically characterized the NFκB p50/RelA heterodimer including the RelA TAD and investigated the TAD’s influence on NFκB-DNA interactions. In solution the RelA TAD is disordered but compact, with helical tendency in two regions that interact with co-activators. The presence of the TAD increased the stoichiometry of NFκB-DNA complexes containing promoter DNA sequences with tandem κB recognition motifs by promoting the binding of NFκB dimers in excess of the number of κB sites. We measured the binding affinity of p50/RelA for DNA containing tandem κB sites and single κB sites. While the presence of the TAD enhanced the binding affinity of p50/RelA for all κB sequences tested, it increased the affinity for non-specific DNA sequences by over 10-fold, leading to an overall decrease in specificity for κB DNA sequences. Our results reveal a novel function of the RelA TAD in promoting binding to non-consensus DNA previously observed by in vivo studies of NFκB-DNA binding in response to strong inflammatory signals.

## INTRODUCTION

Precise control of gene activation and repression is mediated by both protein-protein and protein-DNA interactions involving transcription factor proteins which bind to specific DNA sequences in regulatory regions of genes and control the rate of transcription of mRNA (1). Eukaryotic transcription factors that activate genes often contain a transcription activation domain (TAD), which is a variable domain that binds transcription co-activators, ultimately recruiting the transcription pre-initiation complex and RNA polymerase to initiate transcription (2,3). TADs tend to be intrinsically disordered and enriched in aromatic and acidic amino acids (4-6).

Traditionally, transcription factors were considered modular units with separable DNA binding and transcription activation functions. For this reason, measurements of DNA binding affinity and specificity *in vitro* have often used isolated DNA-binding domains. However, emerging work has shed light on different ways in which TADS and other disordered regions outside the DNA-binding domain can influence transcription factor-DNA interactions. So far, the presence of the TAD generally results in a decrease in DNA binding affinity. For example, intrinsically disordered regions of the *Drosophila melanogaster* Hox protein Ultrabithorax decrease the affinity of the DNA-binding homeodomain for DNA (7), and a disordered region including the TAD of the human transcription factor Forkhead box O4 autoinhibits its DNA binding (8). In at least two cases, TADs can modulate DNA binding specificity. The N-terminal TAD of the transcription factor p53 modulates DNA-binding specificity by contacting the DNA-binding domain and decreasing the binding affinity for non-specific DNA sequences, thereby increasing DNA sequence specificity (9-11). Intrinsically disordered regions outside the DNA-binding domains of the yeast transcription factors Msn2 and Yap1 direct their binding to specific promoter sites *in vivo*, enabling these proteins to differentiate between identical recognition sequences with different flanking regions (12). Given the dearth of quantitative studies of full-length transcription factors *in vitro*, it is unclear whether modulation of DNA binding affinity and/or specificity by disordered domains is a general phenomenon.

Here, we characterize the transcription factor RelA (p65) in its full-length form, including its 230-residue TAD (Fig. 1A). RelA is a member of the NFκB family of transcription factors, which regulates at least 600 genes involved in processes including inflammation, immune response, differentiation, and cell survival (13,14). The NFκB family consists of five proteins, which can form both homo- and heterodimers (15). The most abundant NFκB dimer is the p50/RelA heterodimer, which is the focus of our work here. Both p50 and RelA contain a DNA-binding Rel-homology domain (RHD), which consists of an N-terminal domain (NTD) and a dimerization domain (DD) connected by a short, flexible linker. The inhibitor protein IκBα holds p50/RelA heterodimers in the cytoplasm under resting conditions. In response to stimuli, IκBα is ubiquitinated and degraded, freeing p50/RelA heterodimers to translocate to the nucleus, bind DNA sequences, and regulate transcription (16). To terminate the signal, newly-synthesized IκBα binds to DNA-bound NFκB and facilitates its dissociation from DNA in a process we have called molecular stripping (17,18). A mutant form of IκBα which was defective in stripping NFκB from DNA *in vitro* showed much slower nuclear export of NFκB in cells, demonstrating the physiological function of IκBα-mediated stripping (19).

**Figure 1:**
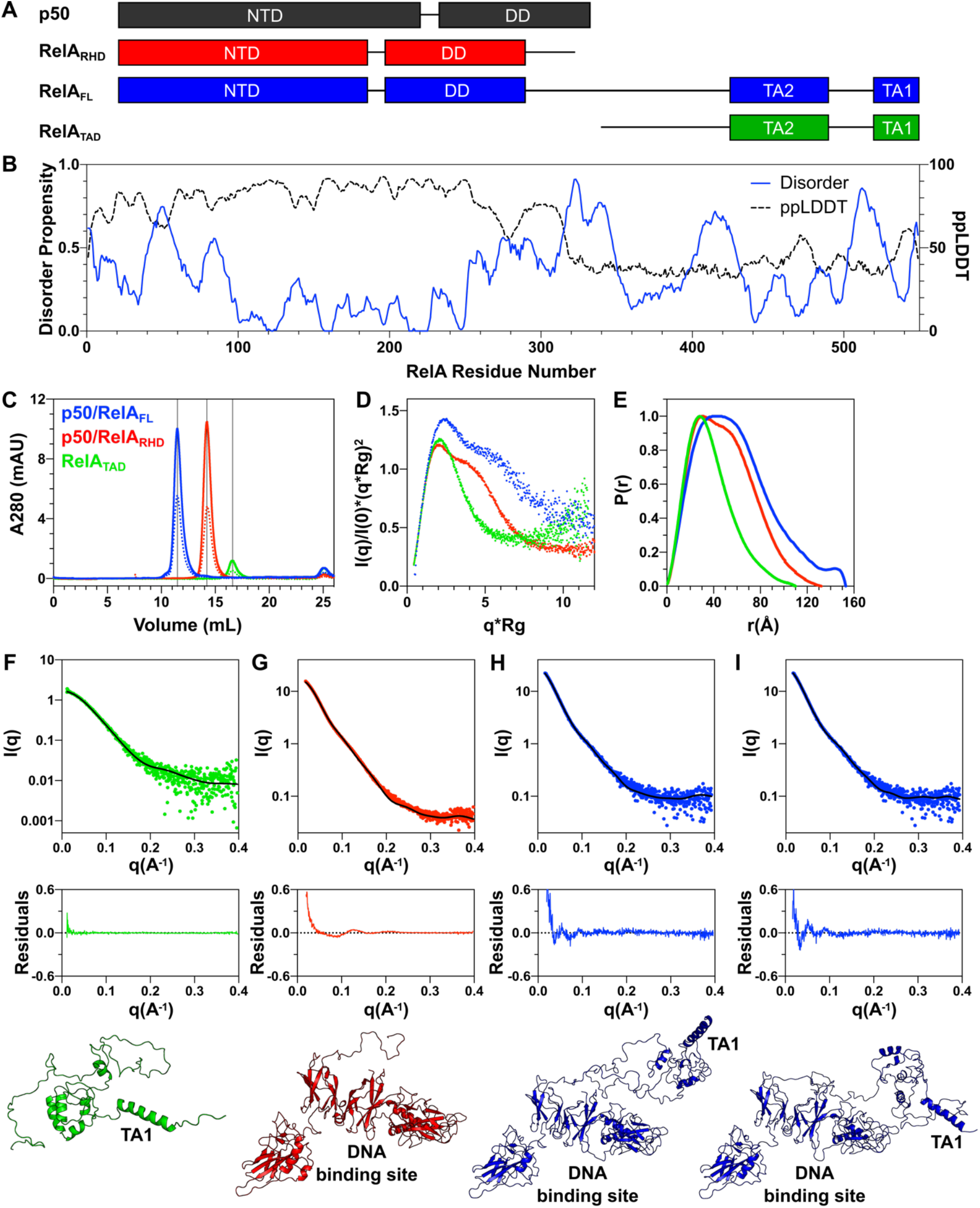
Characterization of NFκB constructs used in this work. A. NFκB subunits p50 and RelA contain well-folded N-terminal domains (NTD) and dimerization domains (DD) that make up the Rel-homology domain (RHD). Additionally, RelA contains an intrinsically disordered transcription activation domain (TAD), which contains two regions important for protein-protein interactions, TA1 and TA2. B. The disorder propensity of full-length RelA (blue, solid) and the predicted AlphaFold2 pLDDT score (black, dashed) were predicted using MetaPredict. The TAD (residues 320-549) is largely predicted to be disordered except for two short stretches corresponding to the TA1 and TA2 motifs. Note that the graph residue numbers are aligned with the RelA_FL_ (residues 19-549) cartoon in panel A. C. The three protein constructs used in this work, p50/RelA_FL_ (blue), p50/RelA_RHD_ (red), and RelA_TAD_ (green) were analyzed by analytical SEC. Solid lines represent 10 µM p50/ RelA_FL_ and p50/RelA_RHD_ and 20 µM RelA_TAD_. Dashed lines represent 5 µM p50/RelA_FL_ and p50/RelA_RHD_ and 10 µM RelA_TAD_. D. SAXS Kratky plots p50/ RelA_FL_ (blue), p50/RelA_RHD_ (red), and RelA_TAD_ (green). E. Pairwise distance distribution for p50/RelA_FL_ (blue), p50/RelA_RHD_ (red), and RelA_TAD_ (green). The R_g_ and D_max_ values of each construct based on SAXS data analysis are in Table 1. F. A model of RelA_TAD_ consistent with the SAXS data was generated using the AWSEM and BilboMD. This model represents one possible conformation of the TAD and reflects its degree of compactness and secondary-structure tendencies in solution. G. A model of p50/RelA_RHD_ consistent with the SAXS data was generated using BilboMD. H and I. Two models of p50/RelA_FL_ were generated using FoxsDock and BilboMD, representing two possible conformations of this protein in good agreement with the SAXS data.

The p50/RelA heterodimer binds to κB DNA sites that contain the consensus sequence GGGRNNYYCC, where R denotes a purine, N denotes any nucleotide, and Y denotes a pyrimidine (20). However, high-throughput *in vitro* analysis has shown that it can bind with similar affinity to some sequences that differ from the consensus, including sequences that contain only a half-site (21,22). Multiple crystal structures have been solved of the p50/RelA heterodimer RHD bound to different DNA sequences, including the immunoglobulin light chain enhancer/HIV long-terminal repeat (LTR) promoter sequence and the IFN-β enhancer sequence (23-26). In all structures, the N-terminal domains of both NFκB subunits contact nucleotide bases within the major groove, whereas the dimerization domains bind each other and interact with the DNA backbone.

**Table 1:** SAXS analysis parameters for p50/RelA_FL_ and p50/RelA_RHD_ and RelA_TAD_.

|  | RelA<sub>TAD</sub> | p50/RelA<sub>RHD</sub> | p50/RelA<sub>FL</sub> |
| --- | --- | --- | --- |
| **R<sub>g</sub> (Å), Guinier Analysis** | 27.3 ± 0.2 | 37.1 ± 0.2 | 46.2 ± 1.3 |
| **R<sub>g</sub> (Å), P(r) analysis** | 27.2 ± 0.1 | 37.1 ± 0.2 | 46.3 ± 0.2 |
| **D<sub>max</sub> (Å), P(r) analysis** | 82 | 110 | 153 |

No structures of NFκB dimers containing both NTDs have been solved in the absence of DNA, and molecular dynamics simulations and single-molecule FRET experiments have demonstrated that the two NTDs in p50/RelA are highly dynamic and can swing apart and create a larger cavity in the absence of DNA (27,28). The RelA TAD is necessary for efficient transcription activation of nearly all target genes (29,30). No structures of the full RelA TAD, either alone or in RelA, are available. Previous structural and biophysical studies have characterized short segments of this domain in isolation, and its behavior within the full-length p50/RelA heterodimer is uncharacterized (31,32).

An important open question in NFκB signaling is how NFκB dimers achieve signaling specificity upon entering the nucleus. As a stress-response transcription factor, NFκB rapidly floods the nucleus in response to stress signals including viral infections and cytokines such as tumor necrosis factor (TNF). In ChIP-Seq experiments, NFκB appears bound non-specifically 15 minutes after TNF treatment in what has been called “global chromatin priming”, whereas stable ChIP-Seq peaks typically arise within 30 minutes (33,34). Nevertheless, across multiple experiments, one third to one half of strong ChIP-Seq peaks bound by RelA did not overlap with a consensus κB motif (35-38). The number of specific κB sites in the human genome was estimated to be ∼10^3^-10^4^, whereas the number of RelA molecules that enter the nucleus in response to a strong activating signal is estimated to be ∼10^5^ (16,39-41). Many questions remain unanswered regarding NFκB specificity, including how *in vivo* results involving full-length RelA relate to *in vitro* experiments lacking the TAD.

Most genes activated by NFκB transcription factors contain multiple κB sites in their promoter and/or enhancer regions, but the function of these multiple sites remains unclear. Tandem DNA binding sites are often associated with cooperative binding and efficient transcription activation (42-44), but the effect of tandem κB sites is not well studied in the context of NFκB-DNA interactions. On a cellular level, many genes activated by NFκB that contain multiple κB sites show a graded response to stimuli, and the transcriptional output by NFκB best matches a model without cooperativity between sites (45). The only DNA sequence containing tandem κB sites that has been studied extensively *in vitro* is the HIV LTR promoter sequence, which has two κB sites separated by four nucleotides. A crystal structure has been solved of the RHDs of two p50/RelA heterodimers bound to the two κB sites, and qualitative experiments have suggested that these two heterodimers may bind with negative cooperativity due to the steric clash generated by the short intervening sequence between the two κB sites (24,46). Binding of NFκB to tandem κB sites has not been studied quantitatively *in vitro*, and no experiments have been performed with an NFκB construct including the RelA TAD.

In this work, we characterize the structural propensity of the RelA TAD alone and in the context of the full-length p50/RelA heterodimer using small-angle x-ray scattering (SAXS), computational modeling, and hydrogen-deuterium exchange mass spectrometry (HDX-MS). We then investigate whether and how the presence of the TAD influences DNA binding stoichiometry, affinity, specificity, and cooperativity to better understand how the full-length p50/RelA heterodimer engages DNA. We find that the RelA TAD is structurally compact but intrinsically disordered both alone and in the context of the p50/RelA heterodimer. Inclusion of the RelA TAD increases the stoichiometry of p50/RelA binding to DNA sequences containing tandem κB sites by promoting binding of p50/RelA dimers in excess of the number of κB sites. It enhances the binding of p50/RelA to all DNA sequences tested, but this effect is more pronounced for DNA sequences that do not match the consensus. Together, these results support a model in which full-length NFκB preferentially binds consensus DNA sequences but also recognizes non-specific DNA, particularly when present in stoichiometric excess as occurs in the nucleus following inflammatory stimulation. This novel role of the RelA TAD helps explain previous *in vivo* observations of widespread non-consensus DNA binding by RelA.

## RESULTS

### Solution characterization of p50/RelA_FL_, p50/RelA_RHD_, and RelA_TAD_

We expressed and purified p50_39-350_/RelA_19-549_ (hereafter referred to as p50/RelA_FL_), p50_39-350_/RelA_19-549_ (hereafter referred to as p50/RelA_RHD_), and RelA_340-549_ (hereafter referred to as RelA_TAD_) protein constructs recombinantly from *E. coli* and characterized the biophysical properties of the RelA TAD alone and within the full-length p50/RelA heterodimer (Fig. 1A). To gain a better understanding of the disordered tendency of the RelA TAD, we used the Metapredict web server (47). Metapredict provides two predicted parameters for a given protein sequence: the disorder propensity, which is designed to reproduce consensus disorder scores from other disorder predictors, and the predicted pLDDT (ppLDDT), which is a prediction of the AlphaFold2 pLDDT (predicted local distance difference test) score. The AlphaFold2 pLDDT score is a residue level confidence metric representing the probability that an AlphaFold structural prediction for a local region will match an experimentally determined structure (48). Low pLDDT scores have been shown to predict disordered regions with high accuracy (49) and provide a useful complement to traditional disorder predictors. The disorder probability scores are low and the ppLDDT scores are high for the first 300 residues of RelA, reflecting the ordered nature of the RHD (Fig. 1B). For the RelA TAD (residues 320-549), the ppLDDT score is below 50% and the disorder propensity score is high with the exception of short stretches including residues ∼430-500 and ∼530-545. These regions correspond to the TA2 and TA1 motifs, respectively, which have previously been shown to form alpha helices in complex with transcription co-activators (31,32). These prediction results paint an overall picture of a largely disordered TAD with local regions of secondary structure propensity.

To confirm that the protein constructs used in this work were soluble and did not form higher-order oligomers, we analyzed the protein samples at two concentrations by analytical size exclusion chromatography (SEC). Each protein eluted as a single symmetric peak from SEC at the expected volume for the mass of the protein construct, and there was no concentration-dependence on elution volume (Fig. 1C). This confirms that the proteins used in this study are soluble and do not form higher-order oligomers at the concentrations used in the experiments presented here.

We used small-angle x-ray scattering (SAXS) to gain insight into the molecular shapes of each protein construct in solution. Kratky analysis of RelA_TAD_ revealed that it adopts a compact structure containing disordered regions, indicated by a Gaussian peak at low q values with a plateau at high q values (Fig. 1D). Kratky analysis of p50/RelA_RHD_ and p50/RelA_FL_ showed both have a multi-domain architecture (Fig.1D), as expected due to the presence of the multidomain RHD. Additionally, the Kratky plot for p50/RelA_FL_ plateaus at high q values, indicating the presence of disordered residues. Average radii of gyration (R_g_) of RelA_TAD_, p50/RelA_RHD_, and p50/RelA_FL_ were determined by Guinier approximation to be 27.3 ± 0.2 Å, 37.1 ± 0.2 Å and 46.2 ± 1.3 Å, respectively (Table 1). Notably, the R_g_ value of 27.3 ± 0.2 Å for RelA_TAD_ is intermediate between the expected R_g_ of a folded protein containing 218 amino acids (∼18-20 Å) and the R_g_ of an excluded-volume polymer of the same size (∼48 Å) (50,51). The P(r) curves of RelA_TAD_, p50/RelA_RHD_, and p50/RelA_FL_ showed D_max_ of 82 Å, 110 Å, 153 Å respectively (Fig. 1E, Table 1). Overall, these results are consistent with the RelA TAD having a compact conformation while retaining a high degree of disorder both when expressed on its own and within p50/RelA_FL_.

To aid in visualization of possible structural conformations, we computationally generated structural models of each construct that were consistent with the SAXS data (Fig. 1F-I, Table 2). Given the dynamic and disordered nature of these protein constructs, our goal was not to produce a single definitive structure but to present models that depict possible configurations of the proteins to enable visualization of their degree of compactness and structural propensity. Models of RelA_TAD_ were initially predicted using the AWSEM code (52) and refined using BilboMD and MultiFoxS to achieve the best agreement with the SAXS data (Fig. 1F, Table 2). The best-fitting representative model has a relatively compact conformation that is mostly unstructured but has some helical content in the TA1 and TA2 regions. We anticipate that the TAD could have both more compact and more extended states within the structural ensemble, and our model represents a conformation with the average R_g_.

**Table 2:** SAXS model parameters for p50/RelA_FL_ and p50/RelA_RHD_ and RelA_TAD_.

|  | RelA<sub>TAD</sub> | p50/RelA<sub>RHD</sub> | p50/RelA<sub>FL</sub><br>Model1/Model2 |
| --- | --- | --- | --- |
| **χ<sup>2</sup>** | 1.1 | 1.35 | 1.44/1.43 |
| **c<sub>1</sub>** | 0.99 | 1.04 | 1.04/1.05 |
| **c<sub>2</sub>** | 3.40 | 0.30 | -0.56/-0.24 |
| **R<sub>g</sub>** | 26.7 | 37.0 | 46.4/45.2 |

A model of p50/RelA_RHD_ consistent with the SAXS data was generated by refining a structure that was generated by removing the DNA from PDB code 1LE5 (25) and running MD simulations for 400 ns. In this structure, the p50 NTD is further away from the RelA NTD, generating a wider DNA-binding cavity (27). This structure was further refined using BilboMD to better fit the SAXS data (Fig. 1G, Table 2). The distance between the two NTDs in this model is wider than in published crystal structures of the NFκB-DNA complex (25) but consistent with single-molecule FRET studies showing the NTDs are dynamic relative to each other and can adopt open conformations in solution (28).

Models of p50/RelA_FL_ were generated by docking the model of RelA_TAD_ onto a model of p50/RelA_RHD_ using FOXSDock. Modeller and BilboMD were used to refine the models that best fit the SAXS data. Excellent χ^2^ values were obtained for two different models in which the TA1 helix extends into solution and the TA2 region is also exposed (Fig. 1H, I, Table 2). The long proline-rich sequence that connects the RHD to the TA2 region adopted several different conformations. These models were neither fully extended nor fully compact, matching the intermediate R_g_ measured by SAXS. We believe the structures represent a subset of the possible conformations of the TAD, which remains highly dynamic in the context of p50/RelA_FL_. Overall, these results match the findings for other acidic, hydrophobic-rich TADs in which hydrophobic and aromatic amino acids important for protein-protein interactions remain solvent-exposed within disordered, negatively charged regions (5,6,53).

### HDX-MS analysis of p50/RelA_FL_ and p50/RelA_RHD_

To gain higher-resolution information about the structure and dynamic properties of p50/RelA_FL_ in solution, we conducted HDX-MS experiments of p50/RelA_FL_ and p50/RelA_RHD_ alone and of each construct of bound to a DNA hairpin containing the HIV-LTR κB sequence. Overall, the results are consistent with the SAXS and modeling data presented above. The models of p50/RelA_FL_ and p50/RelA_RHD_ that best fit the SAXS data did not show stable interactions between the TAD and the RHD. Consistent with these models, no significant differences in deuterium uptake were observed within the RHD whether or not the TAD was present (Fig. 2A). Thus, the RelA TAD does not appear to form long-lived contacts with the RHD that alter the rate of deuterium uptake by this domain. For both protein constructs, peptides contacting the DNA-binding cavity in both the NTDs and DDs of p50 and RelA incorporated significantly less deuterium in the presence of DNA (Fig. 2A, green and yellow peptides). Peptides outside the DNA-binding cavity did not show significant changes in deuterium incorporation upon DNA binding (Fig. 2A, grey and orange peptides). Again, we did not detect significant differences in deuterium incorporation between p50/RelA_FL_ and p50/RelA_RHD_ in the DNA-bound state. Based on these results, we can conclude that the RelA TAD does not form long-lived contacts with the DNA-bound RHD in a manner that alters the rate of deuterium incorporation by the RHD (Fig. 2A).

**Figure 2:**
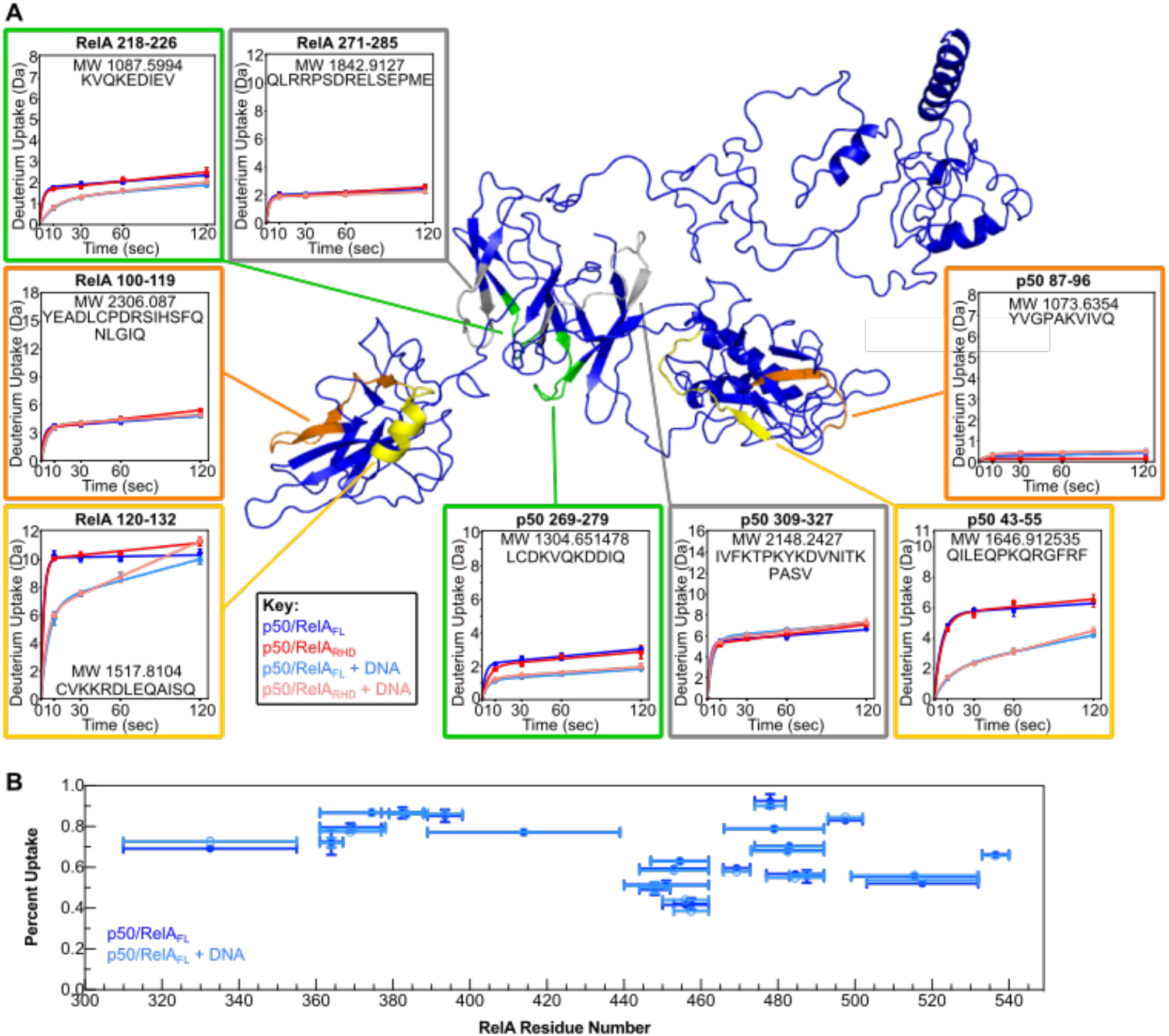
Hydrogen-deuterium exchange analysis of NFκB constructs alone and bound to DNA. A. The structure and dynamics of p50/RelA_FL_ and p50/RelA_RHD_ alone and bound to a DNA hairpin containing the HIV-LTR κB sequence were analyzed using hydrogen-deuterium exchange mass spectrometry (HDX-MS). Representative deuterium uptake plots from peptides from the RHD regions of RelA and p50 are shown mapped to the structural model of p50/RelA_FL_. In general, there were no significant differences in deuterium incorporation when comparing p50/RelA_FL_ vs. p50/RelA_RHD_ alone and p50/RelA_FL_ vs. p50/RelA_RHD_ bound to DNA. Peptides in the DNA-binding cavity showed reduced deuterium incorporation in the presence of DNA (green and yellow peptides highlighted above, representing the DBD and NTD respectively), whereas peptides outside the cavity did not show significant changes in deuterium incorporation in the presence of DNA (grey and orange peptides highlighted above, representing the DBD and NTD respectively). B. The percent deuterium uptake of peptides in the RelA TAD is shown in the presence and absence of DNA. Horizontal bars represent the span of residues in each peptide, and the percent uptake represents the number of deuterons incorporated after 10 seconds relative to the number of exchangeable hydrogen atoms in the peptide. Vertical error bars represent the standard deviation of percent uptake based on three technical replicates.

The RelA TAD showed high levels of deuterium incorporation throughout its sequence, consistent with its disordered, solvent-accessible nature. Most of the TAD had exchanged 70% or more by the 10 second timepoint, with the exception only in the TA1 and TA2 regions (Fig. 2B). Peptides from the regions spanning from residues 440-473, 483-492, and 499-540 showed less than 70% deuterium incorporation at this timepoint, perhaps reflecting their ability to form helical secondary structure as predicted by the AWSEM simulations. Notably, we did not detect differences in deuterium incorporation in peptides from the RelA TAD when comparing the DNA-bound and unbound states (Fig. 2B).

### Binding stoichiometry analysis of NFκB constructs to DNA sequences with tandem sites

To better understand the role of the RelA TAD in binding to DNA, we used electrophoretic mobility shift assays (EMSAs) to investigate the binding of p50/RelA_FL_ and p50/RelA_RHD_ to native DNA promoter sequences containing tandem κB sites. We investigated two tandem sequences: the HIV LTR promoter, which contains two identical κB sites separated by 4 base pairs and the *NFKBIA* promoter, which contains non-identical κB sites separated by 19 base pairs (Fig. 3) (24,54). Notably, the two κB sites in the NFKBIA promoter are quite different from each other, as the second site is a half-site. The HIV LTR promoter has been studied extensively, and a crystal structure of it bound by two p50/RelA RHD dimers has been solved (46). While previous studies have suggested that p50/RelA RHD binds to these sites with negative cooperativity (24,46), the binding affinities driving this association have not been determined quantitatively and has not been investigated in the presence of the RelA TAD. Binding of two NFκB dimers to the *NFKBIA* promoter sequence has not been characterized *in vitro* to our knowledge.

**Figure 3:**
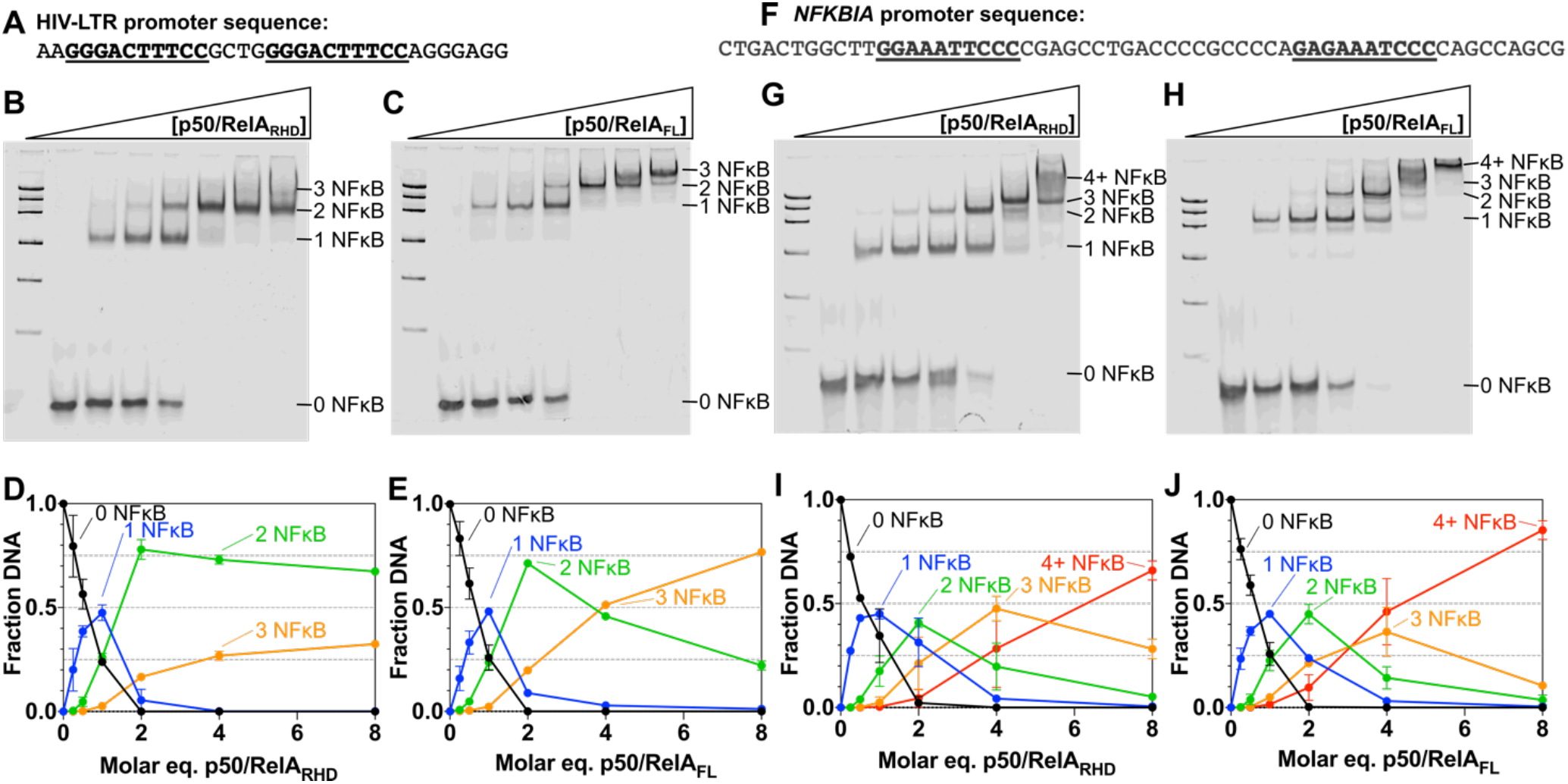
EMSA analysis of binding stoichiometries of p50/RelA_FL_ and p50/RelA_RHD_ to tandem DNA sequences. A. A 33 base pair segment of the HIV LTR promoter sequence was used in these experiments. B. Binding of varying concentrations of p50/RelA_RHD_ (0, 62.5, 125, 250, 500, 1000, and 2000 nM) to the HIV LTR promoter DNA (250 nM) was detected using EMSA. Bands are visible corresponding to free DNA and DNA bound by 1, 2, or 3 p50/RelA_RHD_ dimers. C. Binding of varying concentrations of p50/RelA_FL_ to the HIV LTR promoter DNA (250 nM) was detected using EMSA. The overall patterns are similar to p50/RelA_RHD_, although the band representing DNA bound by three p50/RelA_FL_ dimers is more intense when p50/RelA_FL_ is present in stoichiometric excess (4 or 8 molar equivalents). D and E. The intensities of the bands in the EMSA gels detecting binding of p50/RelA_RHD_ (D) and p50/RelA_FL_ (E) to the HIV LTR DNA were quantified using ImageJ and plotted as a function of NFκB concentration. Data points represent the mean and standard deviation of two independent experiments. F. A 59 base pair segment of the NFKBIA promoter sequence was used in these experiments. G. Binding of varying concentrations p50/RelA_RHD_ (0, 62.5, 125, 250, 500, 1000, and 2000 nM) to the NFKBIA promoter sequence (250 nM) was analyzed using EMSA. Clear bands are present corresponding to free DNA and DNA bound by 1, 2, or 3 p50/RelA_RHD_ dimers, and a smear is present near the top of the gel at higher p50/RelA_RHD_ concentrations, likely representing DNA bound by 4 or more p50/RelA_RHD_ dimers. H. Binding of p50/RelA_FL_ to the NFKBIA promoter sequence was analyzed using EMSA. The overall pattern is similar to the binding of p50/RelA_RHD_, although there is more non-specific binding at high p50/RelA_FL_ concentrations. I and J. The intensities of the bands in the EMSA gels detecting binding of p50/RelA_RHD_ (I) and p50/RelA_FL_ (J) to the NFKBIA DNA were quantified using ImageJ and plotted as a function of NFκB concentration. Data points represent the mean and standard deviation of two independent experiments.

Varying concentrations of NFκB were incubated with the double-stranded DNA segments, then run on a native polyacrylamide gel and stained to observe the distribution of DNA species. For the HIV LTR sequence at low concentrations of NFκB, the EMSA results suggested a lack of binding cooperativity and a similar binding affinity for each κB site. For both p50/RelA_FL_ and p50/RelA_RHD_, the ratio of unbound DNA to DNA bound by a single NFκB dimer to DNA bound by two NFκB dimers is 1:2:1 when NFκB is present at an equimolar concentration to the DNA (Fig 3 B-E). This is the expected distribution of species if NFκB dimers bind the two κB sites in the DNA sequence with the same affinity and without positive or negative cooperativity.

When present in stoichiometric excess, NFκB formed higher-order complexes with the HIV-LTR DNA containing up to three NFκB dimers. This effect was observed for both p50/RelA_FL_ and p50/RelA_RHD_ when present in 8-fold excess but was much more pronounced for p50/RelA_FL_. Binding of more than 2 NFκB dimers to this DNA requires binding to non-consensus sequences. The DNA used in these experiments was 33 base pairs long, sufficiently long to accommodate up to three NFκB dimers, assuming each dimer requires a 10 base pair segment to bind efficiently. However, the DNA does not contain a 10 base pair stretch of non-specific DNA that does not overlap with a κB site. Therefore, it could not accommodate a third NFκB dimer without displacing a dimer bound specifically to a κB site.

The results obtained using the *NFKBIA* promoter DNA are similar in pattern to the HIV-LTR results but more complicated due to the longer length of the *NFKBIA* promoter sequence (59 base pairs). Individual bands can be distinguished for free DNA and DNA bound by 1, 2, or 3 NFκB molecules, and higher-order complexes appear as a smear on the gel. As observed for the HIV-LTR DNA, p50/RelA_FL_ displays a greater propensity toward higher-order complex formation than p50/RelA_RHD_ (Fig. 3 G-J). Importantly, the highest concentration of NFκB used in this experiment (2 µM) and the ratio of NFκB dimers to specific κB sites (4:1), is not unlike the condition in the nucleus, which can be flooded by ∼10^5^ NFκB dimers that recognize ∼10^4^ specific κB sites in response to strong activating signals (41). We estimate the nuclear concentration of p50/RelA heterodimers to be in the 3-4 µM range under such conditions (see Discussion).

We tested the effect of mutating one of the κB sites in the HIV-LTR and *NFKBIA* sequences while leaving the other intact. For HIV-LTR DNA, we found that mutating either the first or the second site resulted in an increase in DNA bound by a single NFκB dimer and a decrease in DNA bound by two NFκB dimers relative to the wild-type DNA sequences when NFκB was present in sub-stoichiometric amounts (Fig. 4 A-D). This result is expected, given that the scrambled DNA sequences contain a single κB site instead of two. Somewhat counterintuitively, at high NFκB concentrations the mutation of one κB site did not diminish NFκB binding and for some sequences actually increased the number of NFκB dimers bound. The amount of DNA bound by three of NFκB dimers when the first site was scrambled was similar to the results obtained using the wild-type DNA (Fig. 3 D and E, Fig, 4 A and B), but we observed significantly more triply-bound DNA when the second site was scrambled (Fig. 4 C and D). This result held for both p50/RelA_RHD_ and p50/RelA_FL_, although again p50/RelA_FL_ was more prone to higher-order complex formation than p50/RelA_RHD_. Notably, scrambling the second site results in 21 sequential base pairs of non-specific DNA which can accommodate two NFκB dimers without displacing the dimer bound to the specific κB site. By contrast, scrambling the first κB site results in only 16 sequential base pairs of non-specific DNA, and in order to bind 3 NFκB dimers this sequence must bind all three non-specifically.

**Figure 4:**
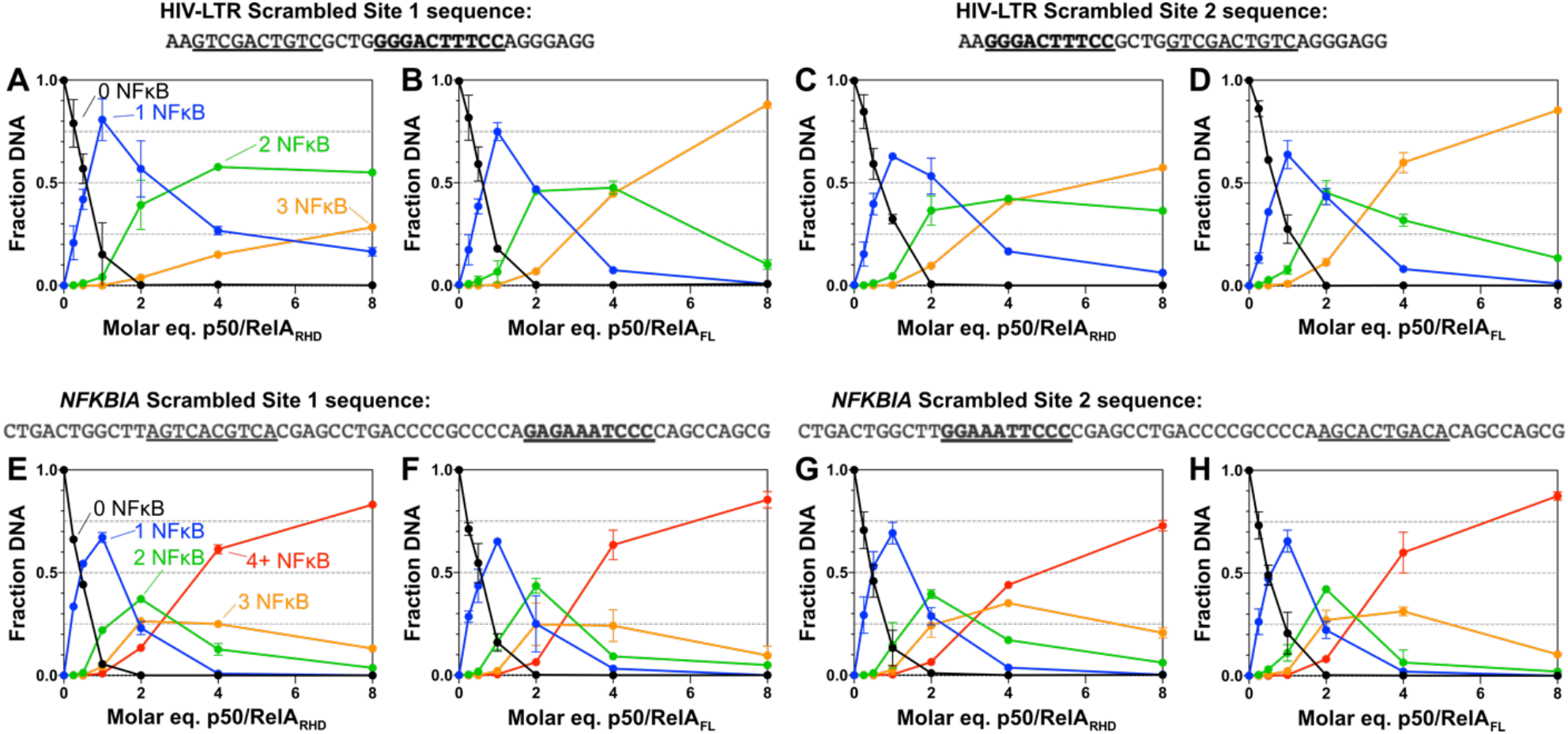
EMSA analysis of binding stoichiometries of p50/RelA_FL_ and p50/RelA_RHD_ to mutated DNA sequences. A and B. Binding of p50/RelA_RHD_ and p50/RelA_FL_ to the HIV LTR sequence in which the first κB site is scrambled was detected using EMSA. Varying concentrations of p50/RelA_RHD_ and p50/RelA_FL_ were mixed with 250 nM HIV-LTR DNA in which the first κB site is scrambled and analyzed using EMSAs. The fraction of DNA bound by 0, 1, 2, or 3 p50/RelA dimers was quantified using ImageJ and plotted as a function of p50/RelA concentration. C and D. When the second κB site in the HIV-LTR promoter DNA segment is scrambled, 21 base pairs of non-specific DNA are present following the first κB site. Varying concentrations of p50/RelA_RHD_ and p50/RelA_FL_ were mixed with 250 nM HIV-LTR DNA in which the second κB site is scrambled and analyzed using EMSAs. Compared to wild-type HIV-LTR DNA and HIV-LTR DNA in which the first site is scrambled, both p50/RelA_RHD_ and p50/RelA_FL_ formed more complexes in which 3 p50/RelA dimers were bound to the DNA. E and F. Similar experiments were conducted using a DNA segment consisting of the *NFKBIA* promoter sequence with the first κB site scrambled. G and H. Binding of p50/RelA_RHD_ and p50/RelA_FL_ to the *NFKBIA* sequence with the second κB site scrambled was also monitored using EMSAs. Overall, scrambling either site in the *NFKBIA* promoter led to an increase in higher-order complex formation at high concentrations of p50/RelA_RHD_ and p50/RelA_FL_. All data points represent the mean and standard deviation of two independent experiments.

The results obtained using the NFKBIA promoter sequence with one site scrambled are generally consistent with the results using the HIV-LTR. For each scrambled NFKBIA sequence, DNA bound by a single NFκB dimer is more prevalent at low concentrations than DNA bound by two NFκB dimers relative to the wild-type DNA, reflecting the presence of only one κB site (Fig. 4 E-H). At higher NFκB concentrations, more complexes formed with 3 or more NFκB molecules bound to the scrambled DNA relative to the wild-type DNA. Again, p50/RelA_FL_ was more prone to higher-order complex formation than p50/RelA_RHD_.

In summary, both p50/RelA_FL_ and p50/RelA_RHD_ can form complexes with DNA in which the number of proteins bound to a strand of DNA exceeds the number of κB sites. For all sequences tested, p50/RelA_FL_ is more prone to formation of these higher-order complexes than p50/RelA_RHD_. Higher-order complexes are more likely to form when stretches of 10 base pairs or longer are accessible without displacement of NFκB dimers from specific κB sites.

### Consequences of specific and non-specific DNA interactions

To better understand how p50/RelA_FL_ might distinguish between specific and non-specific DNA sequences within the nucleus, we conducted EMSA experiments in which we added specific or non-specific hairpin DNA to compete with binding to the HIV-LTR promoter DNA containing tandem κB sites (Fig. 5). Under these conditions, most of the HIV-LTR DNA is bound by 2 p50/RelA_FL_ dimers when p50/RelA_FL_ is present at a 2:1 molar ratio. However, when a DNA hairpin containing the HIV κB sequence is included, it efficiently competes with the HIV-LTR tandem DNA for p50/RelA_FL_ binding. The EMSA band containing HIV-LTR DNA bound by two p50/RelA_FL_ dimers decreases in intensity in the presence of the HIV κB hairpin, while the bands for singly-bound HIV-LTR and free HIV-LTR DNA increase in intensity. A band corresponding to hairpin DNA bound by NFκB is also present.

**Figure 5.**
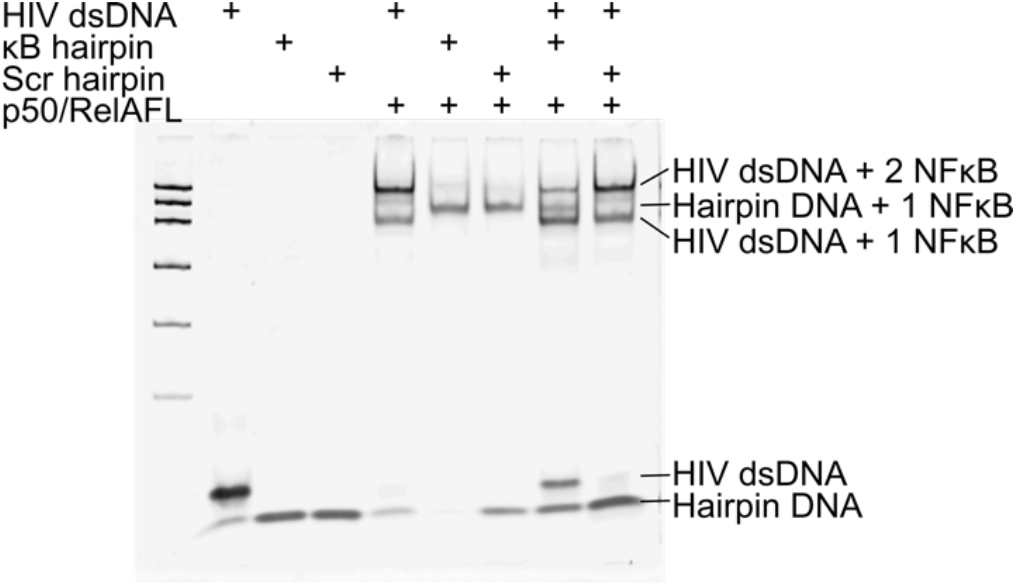
Competition between specific and non-specific DNA sequences for p50/RelA_FL_ binding. 250 nM double-stranded HIV-LTR DNA was incubated with 500 nM p50/RelA_FL_, and hairpin DNA containing either the HIV LTR κB sequence or a scrambled sequence was added to the sample. The κB hairpin was able to efficiently compete with the HIV LTR dsDNA for p50/RelA_FL_ binding (comparing lanes 5 & 8), whereas the scrambled DNA hairpin was unable to compete with the HIV LTR dsDNA for p50/RelA_FL_ binding (comparing lanes 5 & 9).

By contrast, addition of a hairpin with a scrambled DNA sequence does not appreciably change the binding pattern of p50/RelA_FL_ to the HIV-LTR DNA. The p50/RelA_FL_-bound DNA bands match the pattern and intensity of the sample without a competitor hairpin, and a band of free HIV-LTR DNA does not appear (Fig. 5). Therefore, the addition of excess non-specific DNA does not efficiently compete with the specific tandem DNA for binding by NFκB.

We conducted the same experiment using a 59-base pair double-stranded DNA sequence (the *NFKBIA* promoter sequence with both κB sites scrambled) to test whether a longer stretch of non-specific DNA might be a better competitor for NFκB binding. We generally observed higher affinity binding to longer, double-stranded DNA segments than to hairpins in our fluorescence anisotropy assays, so we wanted to test whether excess double-stranded DNA could compete with the specific DNA for NFκB binding. Although these results were harder to interpret due to overlap between NFκB-bound DNA bands, the overall result was the same as with the hairpin DNA. A band of free HIV LTR tandem DNA appeared when the HIV hairpin DNA was included as a competitor but not when the non-specific double-stranded DNA was present (Fig. S1).

Based on these results we believe that despite the lower DNA-binding specificity of p50/RelA_FL_ compared to a construct lacking the TAD, it retains a clear preference for specific κB sites relative to non-specific DNA. Under nucleus-like conditions in which there is an excess of non-specific DNA, we predict it would preferentially bind specific sites. Nevertheless, we anticipate that non-specific DNA interactions may play important roles in NFκB signaling, particularly when NFκB molecules are present in stoichiometric excess relative to the number of κB sites, as is the case during induction of the NFκB signaling system by stressors such as TNF.

### Quantitative determination of DNA binding affinity by p50/RelA_FL_

We used fluorescence anisotropy to determine the equilibrium binding affinities of p50/RelA_FL_ and p50/RelA_RHD_ for several different naturally occurring DNA sequences to gain a more quantitative understanding of how the RelA TAD influences DNA binding. First, DNA hairpins containing the sequences of six κB sites including the NFKBIA promoter, the HIV LTR promoter, the Urokinase promoter, the IFN-β enhancer, and the Rantes promoter were fluorescently labeled on their 5’ ends, and the change in fluorescence anisotropy was monitored upon titration with NFκB constructs (Fig. 6 A-F). In addition, a hairpin containing a random DNA sequence was included to measure binding affinity for non-specific DNA (Fig. 6 G). In general, p50/RelA_FL_ bound to κB DNA sequences with K_d_ values between 1 and 6 nM, whereas p50/RelA_RHD_ bound with a range of K_d_ values between 2 and 20 nM. The p50/RelA_FL_ construct bound with higher affinity to all the DNA sequences tested compared to p50/RelA_RHD_, although the degree to which the inclusion of the TAD enhanced binding affinity varied depending on the sequence. Most dramatically, p50/RelA_FL_ bound the random DNA hairpin with 10-fold higher affinity than p50/RelA_RHD_, 130 ± 40 nM versus 1500 ± 100 nM (Fig. 6 G and H).

**Figure 6:**
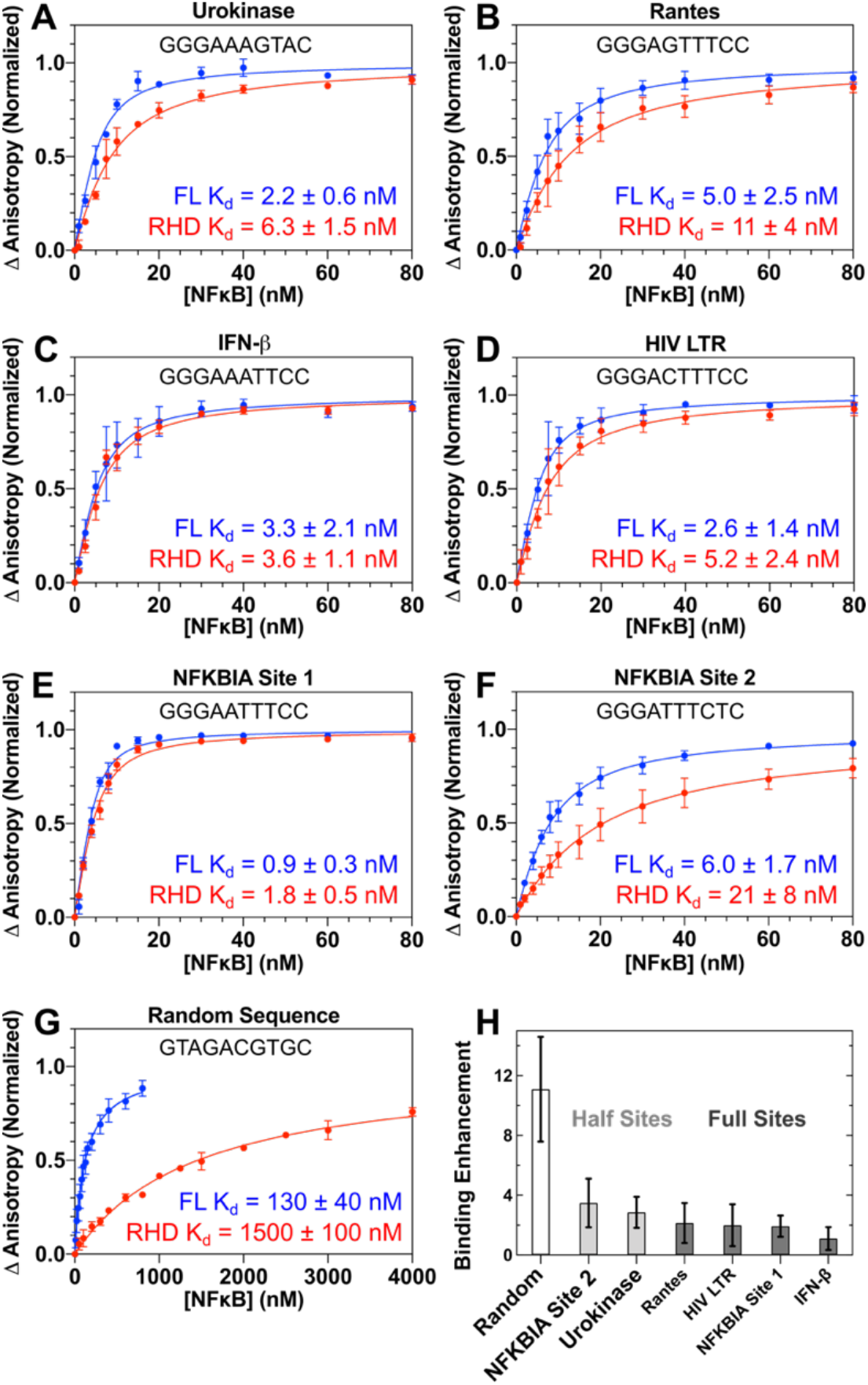
Binding affinities of p50/RelA_FL_ and p50/RelA_RHD_ for hairpin DNA. A-G. DNA hairpins were labeled on the 5’ ends with fluorescein, and 5 nM DNA was incubated with varying concentrations of p50/RelA_FL_ (blue) and p50/RelA_RHD_ (red) to determine binding affinity via fluorescence anisotropy. Data points represent the mean and SEM of three independent experiments, and the reported K_d_ values are the mean and SEM of the K_d_ values derived from each of the experiments. H. The enhancement in binding affinity due to the presence of the TAD in p50/RelA_FL_ was calculated by dividing the K_d_ of p50/RelA_RHD_ by the K_d_ of p50/RelA_FL_ for each DNA sequence. Data points represent mean and SEM. The Urokinase and NFKBIA site 2 sequences are both κB half-sites and show a trend toward slightly greater binding enhancement compared to the full κB sequences.

The binding enhancement determined by fluorescence anisotropy was approximately 3-fold for the hairpins containing κB half-sites (Urokinase and NFKBIA site 2) and 1-2-fold for the hairpins containing full κB sites (Rantes, IFN-β, HIV LTR, and NFKBIA site1). Although the binding enhancements for full versus half site DNA sequences are within error of each other, taken together there is a clear pattern in which the binding enhancement by the TAD is stronger for less-specific DNA sequences. Whereas binding enhancement by the TAD is subtle for full, canonical κB sites, the TAD changes the binding affinity by a full order of magnitude for non-specific DNA. This enhanced binding affinity for non-specific DNA explains the EMSA results showing a greater propensity for p50/RelA_FL_to bind DNA in excess of the number of κB sites (Fig. 3 and 4).

We devised a method to detect binding to each site within the tandem κB sequences individually in order to quantitatively determine the binding affinities for each site within the longer sequence. When a fluorophore is conjugated to the 5’ end of the DNA in close proximity (within 3 base pairs) of a κB site, it becomes immobilized when an NFκB dimer binds to that site, leading to a change in fluorescence anisotropy (55). We found that the change in anisotropy is highly dependent on the distance between the fluorophore and the nearest κB binding site and drops off sharply at distances greater than 3 base pairs (Fig. S2 A-C). By creating double-stranded DNA with 3 base pairs on either side of the tandem κB sites, we can detect binding to either site by monitoring fluorescence anisotropy of a fluorophore conjugated to the 5’ end of either the forward or reverse DNA strand. When the 5’ end of the forward strand is labeled the change in anisotropy reflects binding to the first κB site, whereas when the 5’ end of the reverse strand is labeled the change in anisotropy reflects binding to the second site. Under the conditions used in these experiments, we do not expect binding at the farther site to contribute to the change in anisotropy (Fig. S2 C-D). The results of the binding titrations with labels on each strand were fit globally using MATLAB to determine the binding dissociation constants for each site.

Using this approach, we monitored binding to both κB sites of the HIV LTR sequence and fit the anisotropy data to a simple model in which NFκB bound both sites independently (without positive or negative cooperativity) and with the same affinity (Fig. 7 A). The data for both p50/RelA_FL_ and p50/RelA_RHD_ fit this model well, with K_d_ values of 1.3 ± 0.4 nM and 1.7 ± 0.4 nM, respectively (Fig. 7 B-E).

**Figure 7:**
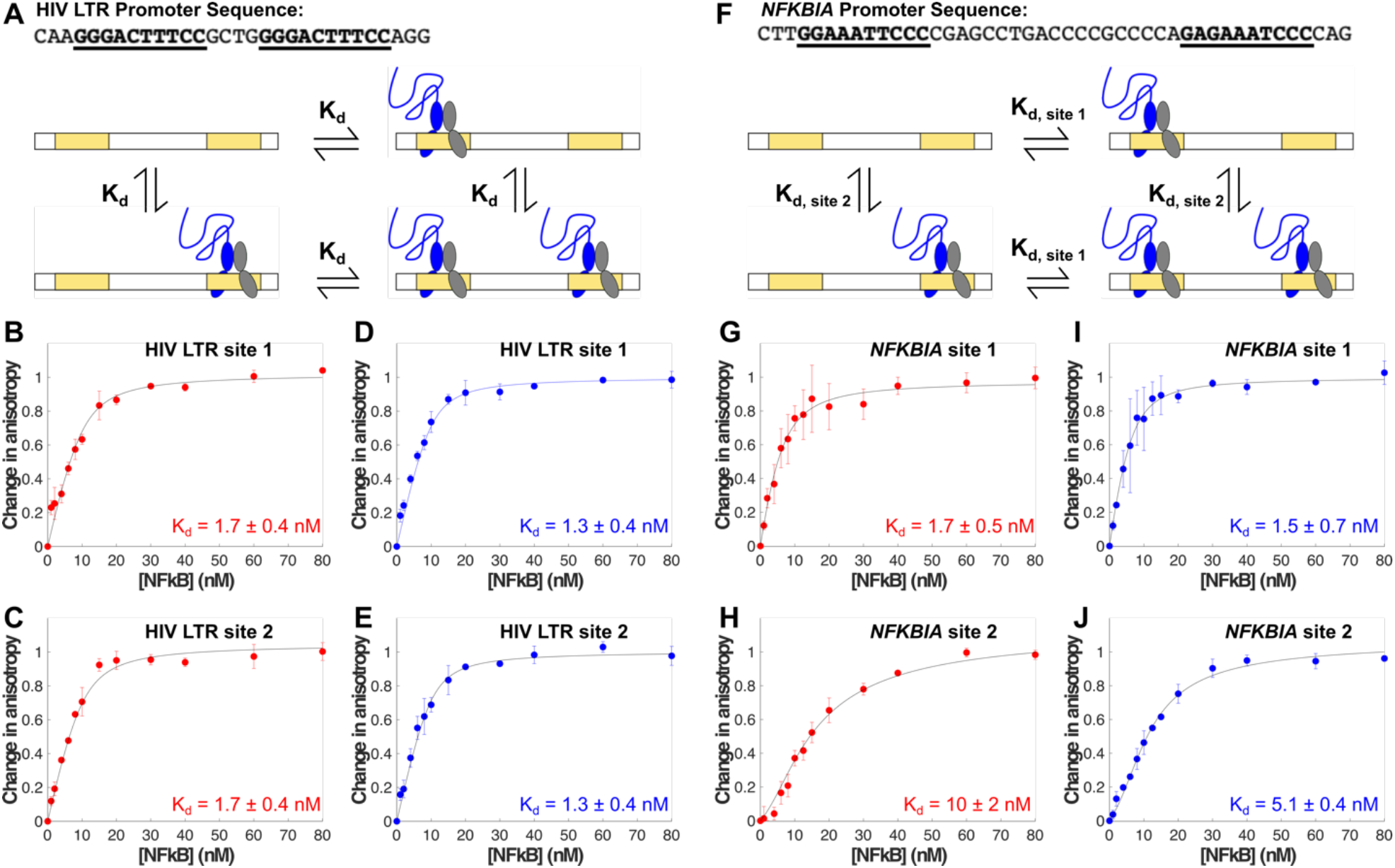
Binding affinities of p50/RelA_FL_ and p50/RelA_RHD_ for κB sites within tandem DNA sequences. A. Double-stranded DNA containing the sequence of the HIV LTR promoter was labeled with fluorescein on the 5’ end of the forward strand to detect binding to the first κB site or the 5’ end of the reverse strand to detect binding to the second κB site. Data obtained for the binding to each site were globally fit to a model in which NFκB binds to each site independently with the same K_d_. B and C. Binding of p50/RelA_RHD_ to the HIV LTR DNA labeled on the 5’ end of forward (B) or reverse (C) strand was detected using fluorescence anisotropy and fit well to a model in which it binds each site independently with a K_d_ of 1.7 ± 0.4 nM. D and E. Binding of p50/RelA_FL_ to the HIV LTR DNA labeled on the 5’ end of forward (D) or reverse (E) strand was detected using fluorescence anisotropy and fit well to a model in which it binds each site independently with a K_d_ of 1.3 ± 0.4 nM. F. Double-stranded DNA containing the sequence of the NFKBIA promoter was labeled with fluorescein on the 5’ end of the forward or reverse strand to detect binding to the first or second κB site, respectively. Fluorescence anisotropy data obtained for binding to each site were globally fit to a model in which NFκB binds to each site independently, with a different K_d_ for each site. G and H. Binding of p50/RelA_RHD_ to the NFKBIA DNA labeled on the 5’ end of forward (G) or reverse (H) strand was detected using fluorescence anisotropy and fit to a model in which it binds each κB site independently, with a K_d_ of 1.7 ± 0.5nM for site 1 and 10 ± 2 nM for site 2. I and J. Binding of p50/RelA_FL_ to the NFKBIA DNA labeled on the 5’ end of forward (I) or reverse (J) strand was detected using fluorescence anisotropy and fit to a model in which it binds each κB site independently, with a K_d_ of 1.5 ± 0.7 nM for site 1 and 5.1 ± 0.4 nM for site 2. Data points shown in this figure are the mean and standard deviation of three independent experiments. K_d_ values are the mean and SEM of the best-fit values determined for each of the three experiments.

We next monitored binding of NFκB to the more complex *NFKBIA* promoter sequence containing tandem κB binding sites. In this case, to obtain a good fit the fluorescence anisotropy data required a more complex model in which the two sites had different K_d_ values but exhibited neither positive nor negative cooperativity (Fig. 7 F). For p50/RelA_FL_, the best-fit K_d_ for site 1 was 1.5 ± 0.7 nM and the best-fit K_d_ for site 2 was 5.1 ± 0.4 nM. For p50/RelA_RHD_, the best-fit K_d_ for site 1 was 1.7 ± 0.5 nM and the best-fit K_d_ for site 2 was 10 ± 2 nM (Fig. 7 G-J). The binding curves for site 2 were sigmoidal in nature, particularly for p50/RelA_RHD_. This reflecting the lower binding affinity for this site and preferential binding to site 1 when the concentration of NFκB is below the concentration of κB binding sites (10 nM).

Overall, the results of these experiments recapitulate the patterns observed using DNA hairpins. For each κB site, the binding of p50/RelA_FL_ is slightly higher affinity than p50/RelA_RHD_. This is especially pronounced for site 2 of the *NFKBIA* promoter, which has a non-canonical half-site sequence. Notably, the data for each tandem sequence fit well to a simple non-cooperative model in which binding to either site is not influenced by binding to the other site. We did not find evidence for either positive or negative cooperativity using this approach, consistent with the EMSA results. Although the binding affinities measured using this approach were generally higher than those measured using hairpin DNA, we believe this is due to the presence of flanking DNA sequences and enhanced stability of longer double-stranded DNA segments rather than a cooperative effect. Importantly, the differences in binding affinities and the relative binding affinities are recapitulated by both the hairpin studies and the studies on tandem sequences.

## DISCUSSION

The results presented here clearly demonstrate that the RelA TAD enhances binding of the p50/RelA heterodimer to nonspecific DNA sequences. This results in higher-order complex formation with long DNA segments containing tandem κB sites and has important implications for our understanding of how the p50/RelA heterodimer engages nuclear DNA and regulates transcription. Multiple studies have found that one third to one half of the genomic sites bound by NFκB do not contain a κB motif (35-40), highlighting the importance and physiological relevance of understanding how NFκB engages both consensus and non-consensus DNA.

The only other protein for which the influence of the intrinsically disordered TAD on DNA binding affinity and specificity has been measured is p53. Our results show that the RelA TAD has the opposite effect of the p53 TAD. First, the presence of the p53 TAD results in a decrease in binding affinity for non-consensus DNA relative to the DNA binding domain alone, whereas the RelA TAD improves binding affinity over that of p50/RelA_RHD_ in all cases. Remarkably, the presence of the RelA TAD dramatically improves binding to non-consensus DNA. Whereas the p53 DNA binding domain binds non-specific DNA with a Kd of 65 nM, the p50/RelA RHD binds non-specific DNA with a K_d_ of 1500 nM (Table 3). The p53 TAD reduces affinity for non-specific DNA by 5.7-fold, whereas the RelA TAD improves binding to non-specific DNA by almost 12-fold (10). This results in the binding affinity of p50/RelA_FL_ for non-specific DNA being even stronger (130 nM) than that for full-length p50 (370 nM).

**Table 3:** K_d_ values of p53* and p50/RelA for consensus and non-consensus DNA

| Protein Construct | Non-consensus DNA $K_d$ | Consensus DNA $K_d$ |
| --- | --- | --- |
| p50/RelA<sub>FL</sub> | $130 \pm 40$ nM | 1-5 nM |
| p50/RelA<sub>RHD</sub> | $1500 \pm 100$ nM | 2-11 nM |
| p53 DBD + TAD | $370 \pm 30$ nM | $16 \pm 2$ nM |
| p53 DBD | $65 \pm 7$ nM | $18 \pm 3$ nM |

\* Values for p53 were obtained from Krois et. al (10).

The cellular concentrations of p50 and RelA have been measured in the ∼200-400 nM range in mouse embryonic fibroblasts and B-cells (56). Given the expected nuclear volume of 10% of the cell volume (57), the concentration of p50/RelA in the nucleus would be 2-4 µM after a strong activating signal. This greatly exceeds the measured binding affinity of 130 nM for p50/RelA_FL_ to non-consensus DNA, and under these conditions nearly all p50/RelA heterodimers would be bound to DNA. Given the presence of only 10^3^-10^4^ consensus κB sites in the human genome, many of the NFκB dimers would be bound to imperfect sites or non-specific DNA regions (39-41). A recent paper that tracked single-molecule transcription factor dynamics in the nucleus found that several different transcription factors had a wide distribution of dwell times, best described by a continuum of affinities for DNA rather than simple specific versus non-specific binding (58). The observation that p50/RelA_FL_ can bind non-specific or imperfect DNA sequences with moderate affinity is consistent with a continuum of binding affinities and likely beneficial to transcription factor function. One possibility is that its binding site search would be more efficient if interactions with non-specific DNA reduced the dimensionality of the search.

Transcription activation by the p50/RelA heterodimer occurs quickly following inflammatory signaling, as NFκB dimers flood the nucleus within the first 15 minutes. Although only weak ChIP-Seq peaks are generally detected within the first 10-15 minutes following inflammatory stimulation, widespread nucleosome repositioning has been observed during this time period prior to the appearance of robust ChIP-Seq peaks at 30 minutes (33,34). Consistent with this *in vivo* result, we recently found that the p50/RelA_RHD_ is capable of binding nucleosomes and unraveling nucleosomal DNA *in vitro* through interactions with both specific κB sequences and nonspecific regions (59). We predict that p50/RelA_FL_ would be even more effective in opening nucleosomes, given its enhanced ability to bind nonspecific DNA sequences. This could have important implications for the search capabilities of p50/RelA, enabling it to more efficiently locate κB sites in both free and nucleosome-bound DNA in order to rapidly activate gene transcription.

The robust ChIP-Seq peaks observed at κB consensus sequences are clearly predicted based on the 1-5 nM binding affinity we measured for p50/RelA_FL_. The ChIP-Seq peaks at non-consensus sequences may arise via a combination of the moderate DNA-binding affinity of ∼100 nM in addition to interactions with other proteins bound at these sites (35,37). A recent study found that multiple human and yeast transcription factors undergo liquid-liquid phase separation mediated by their transcription activation domains *in vitro* when mixed with Mediator subunit MED1 (60). The phenomenon of biomolecular condensation via phase separation has been proposed as a general mechanism for efficient transcription regulation, particularly at nuclear super enhancer sites (61). NFκB has not been shown to undergo liquid-liquid phase separation or biomolecular condensation in the nucleus but is known to associate strongly with super enhancer elements (34). In addition to its involvement in protein-protein interactions that could regulate the formation of large multi-protein assemblies such as super enhancers, the RelA TAD could also promote localization to these elements by facilitating interactions with non-specific DNA within these nuclear sites. Although we did not find evidence for cooperativity between tandem κB motifs, the presence of multiple κB sites within gene regulatory regions could have an avidity effect in localizing NF-κB to these elements without requiring saturation of all κB sites.

## MATERIALS AND METHODS

### Protein expression and purification

N-terminal hexahistidine murine p50_39–350_/RelA_19–321_ (hereafter referred to as p50/RelA_RHD_) was expressed using a modified pET22b vector containing the genes for both polypeptides as described previously (62). The DNA for murine RelA residues 19-549 was synthesized and subcloned into a modified pET22b vector which already contained the gene for N-terminal hexahistidine-p50_39-350_ (hereafter referred to as p50/RelA_FL_). The DNA sequence of the RelA_TAD_ (RelA residues 340-549) was subcloned into pET28a vector with a C-terminal hexahistidine tag.

All vectors were transformed into E. coli BL-21 (DE3) cells and grown to an OD_600_ of 0.5-0.7 at 37°C in M9 minimal media with antibiotic selection. Cultures were cooled on ice for 20 minutes, then protein expression was initiated by the addition of 0.2 mM IPTG. Cultures were incubated at 18°C for 16 hours, then harvested by centrifugation. Pellets were stored at -80°C.

The p50/RelA_RHD_, p50/RelA_FL_, and RelA_TAD_ constructs were lysed by sonication and purified by Ni^2+^-NTA chromatography as described previously for p50/RelA_RHD_ (59). Following overnight dialysis, protein was aliquoted and stored at -80°C. Prior to experiments, aliquots were thawed and further purified. p50/RelA_RHD_ and p50/RelA_FL_ were purified by cation exchange chromatography (MonoS; GE healthcare) to remove bound nucleic acids, as described previously (59). Protein was further purified by size-exclusion chromatography using a Superdex 200 column (GE healthcare) in SEC buffer (25 mM Tris, 150 mM NaCl, 0.5 mM EDTA, 1 mM DTT, adjusted to pH 7.5 at room temperature). Care was taken to separate p50/RelA_FL_ from a breakdown product that eluted at the same volume as p50/RelA_RHD_. RelA_TAD_ was purified by size-exclusion chromatography using a Superdex 75 column, followed by a Superdex 200 column (GE healthcare) in the same buffer.

All purification chromatography steps were conducted in a 4°C cold room. Purity of all proteins was assessed by SDS-PAGE. The protein concentration was determined by absorption at 280 nm using a NanoDrop spectrophotometer. Purified protein was stored at 4°C and all experiments were conducted within 72 hours of purification by size exclusion chromatography.

### Analytical size exclusion chromatography

Protein samples were prepared at two concentrations in SEC buffer. 5 and 10 µM samples of p50/RelA_FL_ and p50/RelA_RHD_ were used, and 10 and 20 µM samples of RelA_TAD_ were used due to its lower molar absorptivity at 280 nm. Samples were injected onto a Superdex 200 10/300 column (GE Life Sciences) equilibrated in the same buffer using a 100 µL sample loop at 4°C.

### Small Angle X-ray scattering (SAXS)

SAXS data were collected at SIBYLS beamline 12.3.1 at the advanced light source (ALS) following standard procedures (63). Three different concentrations (1.25 mg/ml, 2.5 mg/ml 5 mg/ml) of samples (RelA_TAD_, p50/RelA_RHD_, and p50/RelA_FL_) were prepared in SEC buffer. All SAXS data were analyzed either using ATSAS or ScÅtter software. The scattering intensity of the buffer was subtracted from the sample, and the resultant intensity was used for analysis. The radius of gyration (R_g_) was calculated using the Guinier approximation. The dimensionless Kratky plot was generated using the ScÅtter program. The pairwise distance distribution function P(r) was computed using the program GNOM with standard procedures. For the RelA_TAD_ samples, the R_g_ became larger at higher concentrations, so only the lowest concentration (1.25 mg/ml) data were used in the analysis of all three proteins for direct comparisons.

The R_g_ of an excluded-volume polymer containing 218 amino acid residues was calculated to be approximately 48 Å using the equation R_g_ = R_0_*N^ν^, where R_0_ and ν have the values 1.927 Å and 0.598, respectively, as previously determined for denatured proteins in solution (50). The expected R_g_ of a globular protein of this size (18-20 Å) was estimated using the average R_g_ of proteins containing between 201-250 amino acid residues, based on structural characterization of 3412 globular proteins (51).

### Structural modeling based on the SAXS data

To generate a structural model of the RelA TAD (residues 322-549), twenty independent *de novo* AWSEM structure prediction runs were performed to look for folded regions (52). During predictions, the forces guiding folding include backbone terms such as Ramanchandran angle preferences, direct and water-mediated residue-residue contacts, residue burial preferences, hydrogen bonding in α-helices and β-sheets, long-range electrostatic interactions and a bioinformatic local-in-sequence interaction known as the Associative Memory term (64). In each run, the temperature was cooled from 600 K to 100 K over 8 million 2 femtosecond time steps. To generate models of the exact RelA_TAD_ protein construct used for the SAXS data collection, residues 321-339 were removed and a C-terminal 6xHis-tag was added using the web-based server AllosMoD (65) which is integrated with the FoxS server for rigid-body modeling of SAXS data. Next, BilboMD was used to find the conformations of RelA_TAD_ that best matched the SAXS data (66). In BilboMD the helical regions predicted by AWSEM were held stable and the rest of the sequence was allowed to move. We generated 800 conformations at each R_g_ value using the R_g_ determined from the Guinier analysis ± 4 angstroms. The FoXs server was used to calculate the intensity profiles from the RelA_TAD_ conformations generated by BilboMD (67) and each was compared to the SAXS data to determine the best match. The best-fitting model had a χ^2^ value of 1.1.

For p50/RelA_RHD_, we found that the open conformation we previously sampled with atomic MD (27) fit the SAXS data very well. BilboMD was used to further refine the structure using a similar approach as described for RelA_TAD_ and resulted in a model with a χ^2^ value of 1.35.

To generate a model of p50/RelA_FL_ we docked (both PatchDock and FoxDock gave similar results) the RelA_TAD_ model generated by AWSEM simulations onto the model of p50/RelA_RHD_ that best fit the SAXS data. Docking without IκBα present resulted in the TAD binding to the same face of the RHD as IκBα and resulted in models that did not fit the SAXS data well. Therefore, we aligned the dimerization domains of the p50/RelA_RHD_ SAXS model to a crystal structure of the dimerization domains bound by IκBα (PDB 1NFI) (68) and included IκBα as a part of the complex during the docking in order to guide the placement of the TAD away from this site. Several hundred docked models were obtained. The IκBα molecule was removed, and Modeller was used to connect the TAD to the RelA RHD. We used FoXS to calculate the intensity profiles to find the ten models that best agreed with the SAXS data and BilboMD to refine the models. Two models agreed best with χ^2^ of less than 1.4.

### Hydrogen-Deuterium exchange mass spectrometry

Samples containing 5 µM p50/RelA_RHD_ or p50/RelA_FL_ alone or in the presence of 5 µM HIV-LTR hairpin DNA were prepared in SEC buffer. HDX-MS experiments were performed using a Waters Synapt G2Si time-of-flight mass spectrometer equipped with a nanoACUITY UPLC system with HDX technology and a LEAP autosampler. For each time point, 4 µL of sample was incubated at 25°C for 5 minutes, then mixed with 56 µL D_2_O buffer (25 mM Tris, 150 mM NaCl, 1 mM DTT and 0.5 mM EDTA, pH 7.5 at room temperature). Time points were collected in triplicate for 0, 10, 30, 60, and 120 second incubations in D_2_O buffer. Following incubation, 50 µL of the protein solutions was mixed with 60 µL of quench solution (5 M GdnHCl, 0.5% Formic Acid) in a 1°C sample chamber. The pH of this mixture was measured to be 2.7 on ice. The quenched protein solution was then injected onto an in-line pepsin column (immobilized pepsin, Pierce, Inc.). The resulting peptides were trapped and then separated on a C18 column (Acquity UPLC BEH C18, 1.7 µM, 1.0 × 50 mm; Waters Corporation) using a 7-85% acetonitrile gradient with 0.1% formic acid over 7.5 minutes and directly electrosprayed into the mass spectrometer. The mass spectrometer was set to collect data in the Mobility ESI+ mode, with mass acquisition range of 200–2000 (m/z) and scan time of 0.4 s. Continuous lock mass correction was accomplished with an infusion of leu-enkephalin (m/z = 556.277) every 30 s (mass accuracy of 1 ppm for calibration standard). For peptide identification, the mass spectrometer was set to collect data in MS^E^, mobility ESI+ mode instead. The peptides were identified from triplicate MS^E^ analyses of 10 μM of protein solution and data were analyzed using PLGS 2.5 (Waters Corporation). The peptides identified in PLGS were then analyzed in DynamX 3.0 (Waters Corporation). The relative deuterium uptake for each peptide was calculated by comparing the centroids of the mass envelopes of the deuterated samples vs. the undeuterated controls following previously published methods (69). The deuterium uptake was corrected for back-exchange as previously described (70). Deuterium uptake plots were generated in DECA (github.com/komiveslab/DECA) and the data are fitted with an exponential curve for ease of viewing (70).

### Fluorescence anisotropy DNA-binding assay

The DNA oligonucleotides listed below were purchased from Integrated DNA Technologies with a 5’ 5AmMC6 modification. After resuspension in water, approximately 20 nmol DNA was mixed with 300 nmol fluorescein isothiocyanate (Sigma) in a final volume of 100 µL Borax buffer (0.1 M Sodium Tetraborate pH 8.5). Samples were incubated at 70°C for 6 hours, then 500 µL 100% Ethanol was added and samples were stored at -20°C overnight to precipitate the DNA. Precipitated DNA was pelleted by centrifugation, then purified via reverse-phase HPLC as described previously (71). Solvent was removed using a SpeedVac. Hairpin DNA sequences used for the experiments were as follows. Double-stranded DNA sequences are reported in Figures 3 and 4.

IFN-β: 5’ <u>GGGAAATTCC</u>TCCCCCAGGAATTTCCC 3’

Urokinase (UK): 5’ <u>GGGAAAGTAC</u>TCCCCCAGTACTTTCCC 3’

RANTES: 5’ <u>GGGAGTTTCC</u>TCCCCCAGGAAACTCCC 3’

HIV-LTR: 5’ <u>GGGACTTTCC</u>TCCCCCAGGAAAGTCCC 3’

Random: 5’ GTAGACGTGCTCCCCCAGCACGTCTAC 3’

NFKBIA site 1: 5’ TGGAAATTCCCTCCCCCA<u>GGGAATTTCC</u>A 3’

NFKBIA site 2: 5’ AGAGAAATCCCTCCCCCA<u>GGGATTTCTC</u>T 3’

Labeled DNA samples were resuspended in TE buffer, and DNA concentration and labeling efficiency were determined by measuring the absorbance at 260 nm and 495 nm on a NanoDrop spectrophotometer. Double-stranded DNA was created by mixing the labeled strand with an equimolar concentration of the unlabeled complementary strand and incubating in a 95°C heatblock for 5 minutes before turning off the heatblock and allowing the strands to anneal as the sample to slowly cooled to room temperature over the course of 2-3 hours.

Fluorescein-labeled DNA (5 nM) was mixed with varying concentrations of p50/RelA_RHD_ and p50/RelA_FL_ in triplicate in a black 96-well plate and incubated at room temperature for 2.5-3 hours. The fluorescence anisotropy was measured at 22.5 °C on a Beckman Coulter DTX 880 Multimode plate-reader. The excitation and emission wavelengths were 495 nm and 519 nm, respectively. The integration time of the data collection was 1 sec. Anisotropy was calculated using the equation r = [I(V,V) − GI(V,H)]/[I(V,V) − 2GI(V,H)], where r is anisotropy, I(V,V) is the fluorescence intensity in the parallel direction, I(V,H) is the fluorescence intensity in the perpendicular direction and G is the grating factor. The G-factor used for calculating the anisotropy is 0.67, as previously determined for this instrument.

Fluorescence anisotropy data for the hairpin κB DNA sequences were fit to the following equation to determine the equilibrium binding affinity: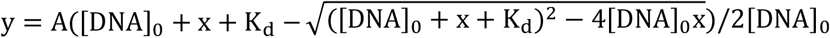, where y is the change in fluorescence anisotropy, A is the maximum change in fluorescence anisotropy, x is the varying concentration of p50/RelA, [DNA]_0_ is the concentration of DNA (5 nM), and K_d_ is the equilibrium binding affinity (72). Anisotropy data for the random hairpin DNA sequence were fit to the equation y = A * x/(x + K).

The fluorescence anisotropy data obtained using DNA sequences with tandem κB sites were fit using an ODE MATLAB-based approach similar to that described previously (73). Detailed equations and approach are described in supplementary material.

DNA binding assays were conducted on three separate days with different DNA and/or protein preparations. For each assay, data were fit to the above equations then normalized by A to determine the fraction bound.

### Electrophoretic mobility shift assays

Samples were prepared in SEC buffer and diluted 1:1 in 2x EMSA loading buffer (40 mM Tris pH 7.5 at room temperature, 100 mM NaCl, 2 mM MgCl_2_, 2 mM DTT, 0.5 mg/mL BSA, 10% glycerol (v/v), and 0.01% bromophenol blue (w/v). Final DNA concentration was 250 nM, and final protein concentrations were 0, 62.5, 125, 250, 500, 1000, and 2000 nM. Samples were incubated for 1 hour at room temperature, then run on 5% polyacrylamide TBE gels in 0.5x TBE buffer in a 4°C cold room. Gels were stained using SYBR Gold nucleic acid stain and imaged using a Typhoon imager. Band intensities were quantified using ImageJ.

## SUPPLEMENTART DATA

SAXS data is available at SASBDB.org; p50/RelARHD heterodimer (SASDHB5), p50/RelAFL (SASDHC5), and RelA Transactivation Domain alone (SASDHD5). These data are available at the following URLs:

https://www.sasbdb.org/data/SASDHB5/3wubsouaf0/ https://www.sasbdb.org/data/SASDHC5/xwlllmre95/ https://www.sasbdb.org/data/SASDHD5/4x8i7qpqzi/

The full project summary is available at https://www.sasbdb.org/project/967/5isz7mjm7q/

## ACKNOWLEDGEMENTS

SAXS data were collected at SIBYLS which is supported by the DOE-BER IDAT and NIGMS ALS-ENABLE (P30 GM124169).

## AUTHOR CONTRIBUTIONS

H.B. designed and performed experiments, analyzed data, and wrote the manuscript. D.N. performed the SAXS experiments. W.C. performed and analyzed simulations. A.V., J.L, M.B. and T.G. performed experiments. P.G.W. supervised simulations. E.A.K. designed experiments and wrote the manuscript.

## SUPPLEMENTARY MATERIAL

### Supplementary Methods

The following ODEs were used to model p50/RelA binding to DNA containing tandem κB sites:

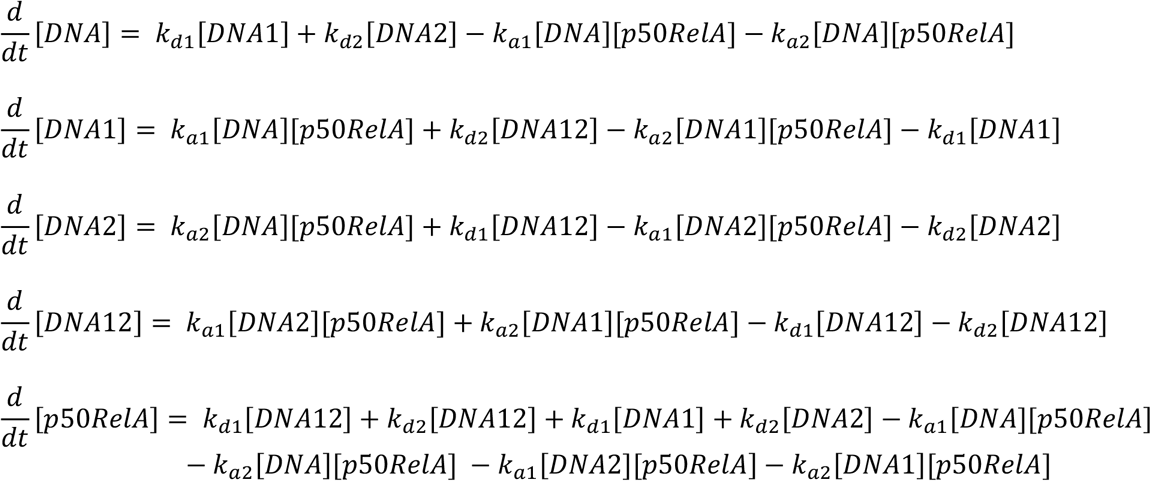

[DNA]: the concentration of free DNA

[DNA1]: the concentration of DNA bound at Site 1 only [DNA2]: the concentration of DNA bound at Site 2 only [DNA12]: the concentration of DNA bound at both sites

[p50RelA]: the concentration of unbound p50/RelA heterodimer k_d1_: the dissociation rate constant of p50/RelA from Site 1

k_d2_: the dissociation rate constant of p50/RelA from Site 2 k_a1_: the association rate constant of p50/RelA to Site 1 k_a2_: the association rate constant of p50/RelA to Site 2

Initial conditions:

[DNA] = 5 nM

[p50RelA] = 0, 1, 2, 4, 6, 8, 10, 15, 20, 30, 40, 60, 80 nM [DNA1] = [DNA2] = [DNA12] = 0 nM

k_a1_ = k_a2_ = 10_9_ M_-1_s_-1_

k_d1_ and k_d2_ were fit to determine the best-fitting K_d_ values for sites 1 and 2 (K_d_ = k_d_ / k_a_). For HIV LTR DNA, k_d1_ = k_d2_.

The measured change in fluorescence anisotropy reflects the fraction of the DNA bound at site 1 or site 2 when the label is placed on the 5’ end of the forward or reverse strand, respectively.

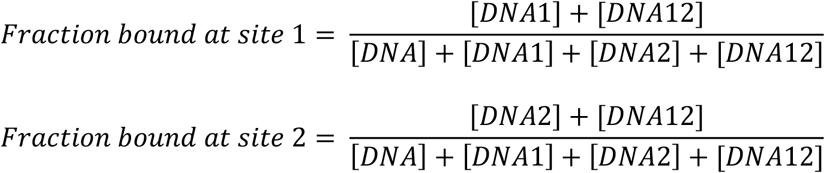

**Supplementary Figure 1:**
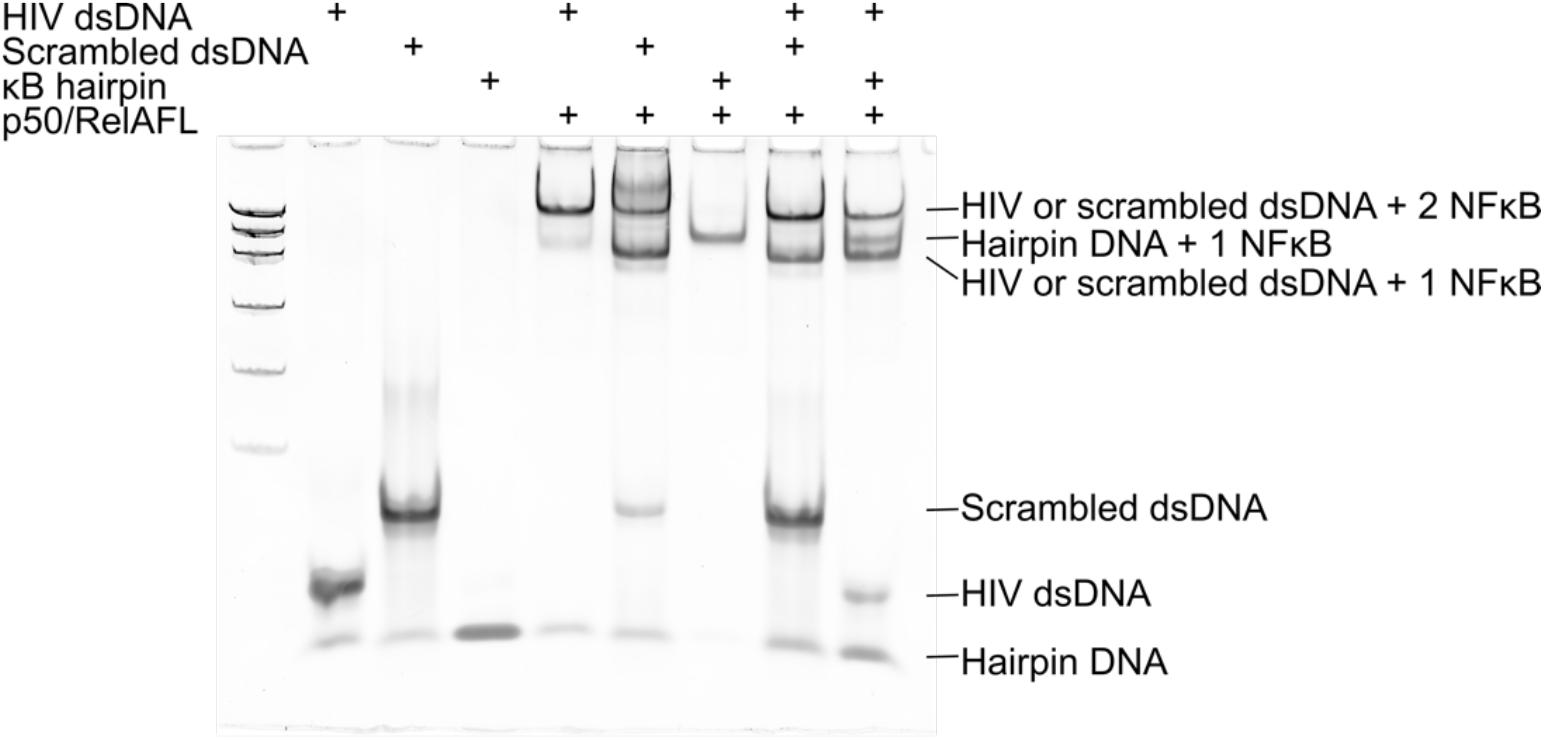
Competition between specific and non-specific DNA sequences for p50/RelA_FL_ binding. An EMSA experiment was conducted to test the ability of specific and non-specific DNA sequences to compete with the HIV LTR sequence for p50/RelA_FL_ binding. 250 nM double-stranded HIV-LTR DNA was incubated with 500 nM p50/RelA_FL_, and 250 hairpin DNA containing either the HIV LTR κB sequence or double-stranded DNA containing the NFKBIA sequence with both κB sites scrambled were added to the sample. The κB hairpin was able to efficiently compete with the HIV LTR dsDNA for p50/RelA_FL_ binding (comparing lanes 5 & 9). The top band, corresponding to HIV LTR dsDNA bound by two p50/RelA_FL_ dimers, decreases in intensity, whereas a band corresponding to the hairpin DNA bound by a p50/RelA_FL_ dimer and a band corresponding to free HIV LTR dsDNA both appear. By contrast, the scrambled dsDNA does not efficiently compete with the HIV LTR dsDNA for p50/RelA_FL_ binding (comparing lanes 5 & 8). The band corresponding to free HIV LTR dsDNA does not appear in this lane, whereas the band corresponding to free scrambled dsDNA remains strong.

**Supplementary Figure 2:**
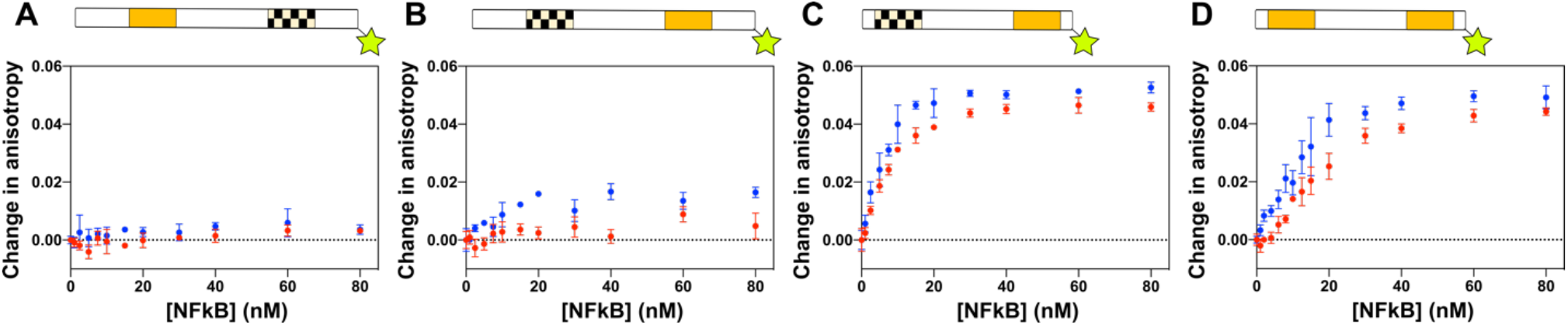
Dependence of fluorescence anisotropy change on fluorophore placement. **A**. When the fluorophore is conjugated 38 base pairs from a κB site, there is no observed change in fluorescence anisotropy as a function of NFκB concentration. The DNA sequence used in this experiment is the *NFKBIA* promoter with the second site scrambled (see Fig. 6G-H). **B**. When the fluorophore is 9 base pairs away from a κB site, there is only a slight increase in fluorescence anisotropy upon titration with NFκB. The DNA sequence used here is the *NFKBIA* promoter with the first site scrambled (see Fig. 6 E-F). **C**. When the distance between the fluorophore and the κB site is reduced to 3 base pairs, titration with NFκB results in a much greater change in anisotropy. In this experiment, the first κB site of the *NFKBIA* sequence is scrambled so the change in anisotropy reflects binding to only the second site. **D**. When both κB sites are intact in the *NFKBIA* promoter, the fluorescence anisotropy change is the same as in panel C, which uses the same DNA sequence but with only one intact κB site. Therefore, the observed anisotropy change can all be accounted for by binding interactions with the κB site nearest the fluorophore. All data points represent the mean and standard deviation of three technical replicates. Checked boxes represent scrambled κB sites, and yellow boxes represent intact κB sites. Blue points are for p50/RelA_FL_ and red points are for p50/RelA_RHD_.

