## Supplemental Materials for "An intrinsically disordered transcription activation domain alters the DNA binding affinity and specificity of NFκB p50/RelA"

### Supplementary Material

#### Supplementary Methods:

The following ODEs were used to model p50/RelA binding to DNA containing tandem κB sites:

$$\frac{d}{dt}[DNA] = k_{d1}[DNA1] + k_{d2}[DNA2] - k_{a1}[DNA][p50RelA] - k_{a2}[DNA][p50RelA]$$

$$\frac{d}{dt}[DNA1] = k_{a1}[DNA][p50RelA] + k_{d2}[DNA12] - k_{a2}[DNA1][p50RelA] - k_{d1}[DNA1]$$

$$\frac{d}{dt}[DNA2] = k_{a2}[DNA][p50RelA] + k_{d1}[DNA12] - k_{a1}[DNA2][p50RelA] - k_{d2}[DNA2]$$

$$\frac{d}{dt}[DNA12] = k_{a1}[DNA2][p50RelA] + k_{a2}[DNA1][p50RelA] - k_{d1}[DNA12] - k_{d2}[DNA12]$$

$$\frac{d}{dt}[p50RelA] = k_{d1}[DNA12] + k_{d2}[DNA12] + k_{d1}[DNA1] + k_{d2}[DNA2] - k_{a1}[DNA][p50RelA] - k_{a2}[DNA][p50RelA] - k_{a1}[DNA2][p50RelA] - k_{a2}[DNA1][p50RelA]$$

[DNA]: the concentration of free DNA
[DNA1]: the concentration of DNA bound at Site 1 only
[DNA2]: the concentration of DNA bound at Site 2 only
[DNA12]: the concentration of DNA bound at both sites
[p50RelA]: the concentration of unbound p50/RelA heterodimer
$k_{d1}$: the dissociation rate constant of p50/RelA from Site 1
$k_{d2}$: the dissociation rate constant of p50/RelA from Site 2
$k_{a1}$: the association rate constant of p50/RelA to Site 1
$k_{a2}$: the association rate constant of p50/RelA to Site 2

Initial conditions:
[DNA] = 5 nM
[p50RelA] = 0, 1, 2, 4, 6, 8, 10, 15, 20, 30, 40, 60, 80 nM
[DNA1] = [DNA2] = [DNA12] = 0 nM
$k_{a1} = k_{a2} = 10^9\ M^{-1}s^{-1}$

$k_{d1}$ and $k_{d2}$ were fit to determine the best-fitting $K_d$ values for sites 1 and 2 ($K_d = k_d / k_a$).
For HIV LTR DNA, $k_{d1} = k_{d2}$.

The measured change in fluorescence anisotropy reflects the fraction of the DNA bound at site 1 or site 2 when the label is placed on the 5' end of the forward or reverse strand, respectively.

$$Fraction\ bound\ at\ site\ 1 = \frac{[DNA1] + [DNA12]}{[DNA] + [DNA1] + [DNA2] + [DNA12]}$$

$$Fraction\ bound\ at\ site\ 2 = \frac{[DNA2] + [DNA12]}{[DNA] + [DNA1] + [DNA2] + [DNA12]}$$

### Supplementary Figure 1:

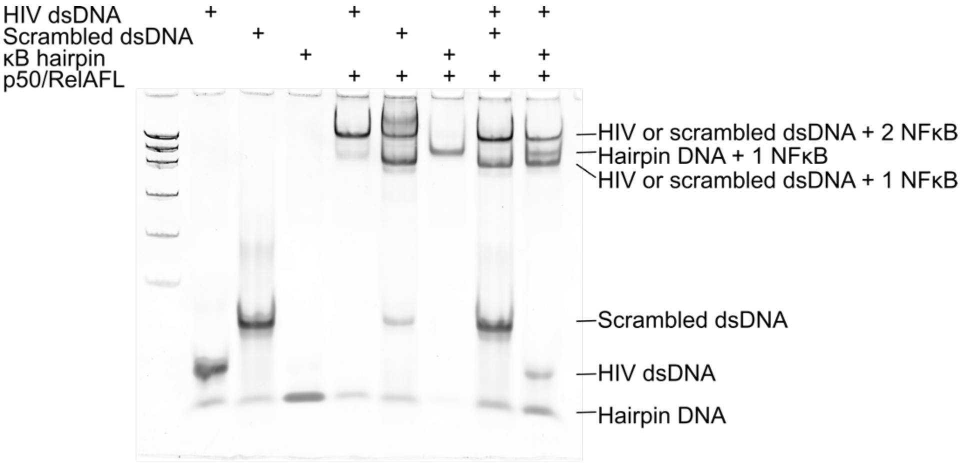


Competition between specific and non-specific DNA sequences for p50/RelA$_{FL}$ binding. An EMSA experiment was conducted to test the ability of specific and non-specific DNA sequences to compete with the HIV LTR sequence for p50/RelA$_{FL}$ binding. 250 nM double-stranded HIV-LTR DNA was incubated with 500 nM p50/RelA$_{FL}$, and 250 hairpin DNA containing either the HIV LTR κB sequence or double-stranded DNA containing the NFKBIA sequence with both κB sites scrambled were added to the sample. The κB hairpin was able to efficiently compete with the HIV LTR dsDNA for p50/RelA$_{FL}$ binding (comparing lanes 5 & 9). The top band, corresponding to HIV LTR dsDNA bound by two p50/RelA$_{FL}$ dimers, decreases in intensity, whereas a band corresponding to the hairpin DNA bound by a p50/RelA$_{FL}$ dimer and a band corresponding to free HIV LTR dsDNA both appear. By contrast, the scrambled dsDNA does not efficiently compete with the HIV LTR dsDNA for p50/RelA$_{FL}$ binding (comparing lanes 5 & 8). The band corresponding to free HIV LTR dsDNA does not appear in this lane, whereas the band corresponding to free scrambled dsDNA remains strong.

### Supplementary Figure 2:

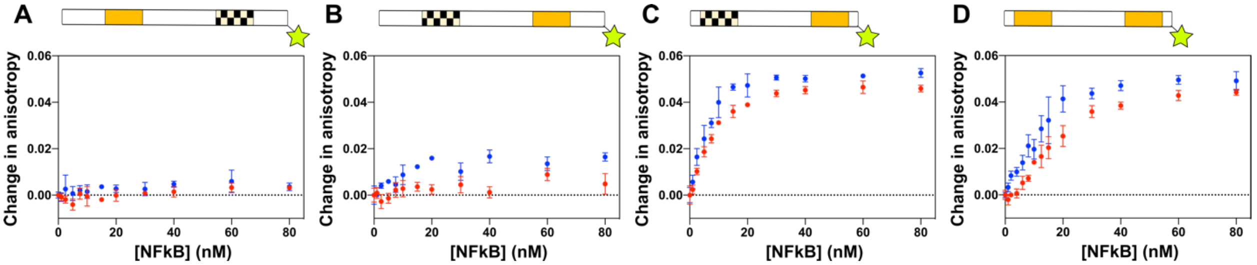


Dependence of fluorescence anisotropy change on fluorophore placement. **A.** When the fluorophore is conjugated 38 base pairs from a κB site, there is no observed change in fluorescence anisotropy as a function of NFκB concentration. The DNA sequence used in this experiment is the *NFKBIA* promoter with the second site scrambled (see Fig. 6G-H). **B.** When the fluorophore is 9 base pairs away from a κB site, there is only a slight increase in fluorescence anisotropy upon titration with NFκB. The DNA sequence used here is the *NFKBIA* promoter with the first site scrambled (see Fig. 6 E-F). **C.** When the distance between the fluorophore and the κB site is reduced to 3 base pairs, titration with NFκB results in a much greater change in anisotropy. In this experiment, the first κB site of the *NFKBIA* sequence is scrambled so the change in anisotropy reflects binding to only the second site. **D.** When both κB sites are intact in the *NFKBIA* promoter, the fluorescence anisotropy change is the same as in panel C, which uses the same DNA sequence but with only one intact κB site. Therefore, the observed anisotropy change can all be accounted for by binding interactions with the κB site nearest the fluorophore. All data points represent the mean and standard deviation of three technical replicates. Checked boxes represent scrambled κB sites, and yellow boxes represent intact κB sites. Blue points are for p50/RelA$_{FL}$ and red points are for p50/RelA$_{RHD}$.
